# Species richness and community structure of bats along a forest elevational transect in Papua New Guinea

**DOI:** 10.1101/2022.02.17.480839

**Authors:** Elise Sivault, Pita K. Amick, Kyle N. Armstrong, Vojtech Novotny, Katerina Sam

**Affiliations:** Faculty of Science, University of South Bohemia, Branisovska 1760, 370 05 Ceske Budejovice, Czech Republic; Biology Centre of the Czech Academy of Sciences, Institute of Entomology, Branisovska 31, 370 05 Ceske Budejovice, Czech Republic; Biological Science Division, University of Papua New Guinea, PO Box 320, Papua New Guinea; The New Guinea Binatang Research Centre, PO Box 604, Madang, Papua New Guinea; Amick Environmental Consulting, PO Box 1179 Mt Hagen, 511 Western Highlands Province, Papua New Guinea; University of Adelaide, North Terrace, Adelaide, South Australia 5005; South Australian Museum, North Terrace, Adelaide, South Australia 5000

**Keywords:** bat communities, species richness, tropical montane, elevational gradient, Papua New Guinea

## Abstract

Over the past decades, elevational gradients have become a powerful tool with which to understand the underlying cause(s) of biodiversity. The Mt. Wilhelm elevational transect is one such example, having been used to study the birds, insects, and plants of Papua New Guinea (PNG). However, a survey of mammals from this forest elevational transect was lacking. We thus aimed to investigate patterns in the community structure and species richness of bats (Chiroptera) along the transect, link the species to available regional data, and explain the observed patterns by including environmental characteristics. Bat communities were surveyed between 200 m and a timberline at 3,700 m a.s.l. at eight study sites separated by 500 m in elevation. We conducted mist-netting and acoustic surveys to detect and identify species at each site. Regional data were compiled to compare local with regional diversity. Finally, biotic (i.e., food availability, habitat features) and abiotic (i.e., mean daily temperature, available land area) factors were included in our analyses to disentangle the ecological drivers underlying bat diversity. Results revealed that species richness decreases with ascending elevation and was best explained by a corresponding decrease in both area and temperature. We also observed community turnover along the transect at local and regional scales, along with the increase of species’ elevational ranges. Consequently, despite that the study was restricted to one mountain in PNG, it demonstrates how basic inventory surveys can be used to address ecological questions in other similar and undisturbed tropical mountains.

## 1. INTRODUCTION

Mountains are considered important biodiversity hotspots due to the great richness of endemic species that occur on them (Gradstein et al., 2008; Noroozi et al., 2018). They are also one of the most anthropogenically threatened environments in the world (Davis & Shaw, 2001; Ricketts et al., 2005). Consequently, elevational gradients are excellent systems for the study of biodiversity, global change, and conservation perspectives. Often used in models as a proxy for climate change, they allow us to study animal and plant responses to changes in biotic and abiotic factors (McCain & Colwell, 2011). Additionally, they can reflect responses to land-use changes, which often occur at low elevations (Becker et al., 2007). For these reasons, community studies, and especially of patterns of species richness along elevational gradients, have remained popular for many decades (Stevens et al., 2019). Several meta-analyses of terrestrial vertebrate groups have emerged in recent times that demonstrate varying trends in species diversity according to geographical location, largely because of their significant climatic differences (i.e., temperature, humidity) (e.g., McCain, 2005, 2007b, 2009, 2010).

Among terrestrial vertebrate groups, bats are considered commonly in studies of temperate (Piksa et al., 2013; Scherrer et al., 2019), South American, and African tropical mountain ecosystems (Carvalho et al., 2019; Mongombe et al., 2019; Reardon & Schoeman, 2017). Two predominant patterns of bat species richness have been observed—a decrease with elevation in most tropical regions, and a unimodal trend in temperate regions (McCain, 2007b; McCain & Grytnes, 2010). The area hypothesis states that the amount of land area for each elevational band (e.g., 100–200m) on a mountain will be positively related to the diversity observed in that band (Terborgh, 1973), however, bats showed either no significant relationship or a negative association between species richness and available area (McCain, 2007a). It has been suggested that the highest bat species richness occurs in the elevational zone where water and temperature are simultaneously high (i.e., low and mid-elevations in tropical and temperate mountains respectively) (McCain, 2007b). Being small and volant, bats spend much of their energy budget on flight and thermoregulation, which is dependent on ambient temperature and therefore, limits their distribution in cold temperature regimes (Graham, 1983; McNab, 1982). In addition, water availability and temperature indirectly influence food resource availability (e.g., through fruiting tree phenology, the abundance of arthropods), and vegetation (e.g., shrub density), thereby influencing foraging behaviour and the availability of roosting sites (Charbonnier et al., 2016; Moura et al., 2016). While abiotic factors (i.e., temperature, available area within an elevation band) have been explored often in existing models, biotic factors as food resources and habitat characteristics have been considered rarely.

In addition to species richness, the species composition of bat assemblages can vary along an elevational gradient under different scenarios. It has been suggested that high-elevation bat species are able to exist at all elevations because of broad physiological tolerance and ecological requirements (e.g., Stevens, 1992). However, under a scenario of climate change, bat species are acclimated to high mountain conditions because they are geographically, ecologically, and/or physiologically constrained to high elevations (e.g., LaVal, 2004). Thus, species found only at high elevations are likely to expand their ranges or be strictly constrained by recent events of climate change.

Feeding specialization might be another factor affecting assemblage composition. Despite that bat communities are typically dominated by insectivores, the relative species richness of frugivorous and nectarivorous bats peaks in the tropics, especially in the Neotropics, Oceania, and Australasia (Maas et al., 2016). However, patterns in the distribution of bat specialization along elevational gradients have rarely been documented, despite plant and insect distributions varying greatly. Fruiting trees are typically reported to decline in diversity and abundance with increasing elevation (Loiselle & Blake, 1991) while insects follow various patterns (i.e., none, peaking mid-elevation, decreasing, increasing) according to their group and/or localities (Hodkinson, 2005). Consequently, elevation might act as a filter of bat feeding guilds and impact the species composition of bat assemblages.

PNG bats represent seven percent of the world’s bat diversity (Bonaccorso, 1998; Mammal Diversity Database, 2021). From a total of 95 native species, PNG has at least 19 endemic bat species (Bonaccorso, 1998). In recent decades, this unique richness attracted new research focused on viruses (Breed et al., 2010; Field et al., 2013), metabolism (McNab & Bonaccorso, 2001), and the home range of single species (Bonaccorso et al., 2002; Winkelmann et al., 2000, 2003). However, there is a lack of knowledge of bat community structure due to limited specific focus within larger mammal studies (Helgen, 2007; Helgen et al., 2011). Much of the effort for bat research in the past two decades has been as part of basic inventory surveys, environmental impact assessments and monitoring for industry (Armstrong et al., 2020; Kale et al., 2018; K.P. Aplin and K.N. Armstrong unpublished reports), university research (Bonaccorso et al., 2002; Robson et al., 2012; Wiantoro, 2020) or else as part of biodiversity assessments for conservation organizations (Armstrong et al., 2015a,b; Armstrong & Aplin, 2011, 2014). While there is no central library of PNG bat echolocation calls, these studies have steadily accumulated knowledge and resources that support both acoustics-based and genetics-based identification, underpin recent species profile revisions in the IUCN Red List, and studies of taxonomic resolution (e.g., Wiantoro, 2020). They also mark a shift towards a primary reliance on acoustics- based detection and identification on field surveys, rather than trapping as was relied upon in the past, though trapping is still the best means of surveying for small species in the Pteropodidae and collecting material to allow robust acoustics-based identification.

As a part of the Bismarck Range, Mt. Wilhelm is the highest peak in PNG (4,509 m a.s.l.) and offers a complete elevational transect in relatively intact tropical forests. Established study transects have become well-studied for birds (Marki et al., 2016; Sam et al., 2017, 2019), insects (Cesne et al., 2015; Finnie et al., 2021; Novotny et al., 2005; Orivel et al., 2018; Souto_Vilarós et al., 2020; Szczepański et al., 2018), and plants (Lofthus et al., 2020; Smith, 1977; Volf et al., 2020). Thus, given what previous efforts on other biota offer, the Mt. Wilhelm transect provided an opportunity not found elsewhere in New Guinea to study bat communities. Consequently, the present study aimed to: (a) document bat species richness patterns and community assemblages with increasing elevation to determine whether elevation is a filter of specific bat species and/or feeding guilds. We expected to see a steeply decreasing pattern in species richness, as it has been typically observed on tropical wet mountains; (b) investigate which of the abiotic (i.e., mean daily temperature, available land area) and biotic (i.e., habitat, food availability) factors drive bat diversity patterns. We assumed that temperature will best explain the patterns as described in the majority of past studies; (c) compare bat communities from the Mt. Wilhelm transect with the regional data (compiled from Bonaccorso, 1998), as well as the bat elevation ranges to determine whether some species are out of their previously recorded ranges. We expected that species we detect in highlands will have wider environmental tolerance, thus they will occur also in the lowlands, nevertheless, following the current scenario of climate change, we also expected to find species strictly constraint to high elevations.

## 2. METHODS

### 2.1 Study area

We surveyed bat communities along the elevational transect of Mt. Wilhelm in PNG between 200 m and 3,700 m a.s.l. at eight elevational study sites separated by 500 m elevational increments (i.e., 200 m, 700 m, 1,200 m, 1,700 m, 2,200 m, 2,700 m, 3,200 m, and 3,700 m a.s.l. and ±60 m for each study site due to the rough terrain). The 30 km long elevational transect, which stretches between 5°44’S, 145°2’E and 5°47’S, 145°03’E, is located along the Bismarck Range’s northern slope (Figure 1). Vegetation types used here follow Paijmans (1975), i.e., lowland alluvial forest (<500 m a.s.l.), foothill forest (501–1,500 m a.s.l.), lower montane forest (1,501–3,000 m a.s.l.), and upper montane forest (>3,000 m a.s.l.) (Figure S1.1 in Appendix S1 in supporting information). Mean daily temperature decreases linearly (r = - 0.9) from 27.4°C at the 200 m a.s.l to 8.37°C at the timberline (3,700 m a.s.l) (Table S3.2) (Sam et al., 2019). The average annual precipitation is 3,288 mm (local meteorological station) in the lowlands, rising to 4,400 mm at the forest edge, with a distinct condensation zone between 2,500 and 2,700 m a.s.l. (Sam et al., 2019).

**Figure 1:**
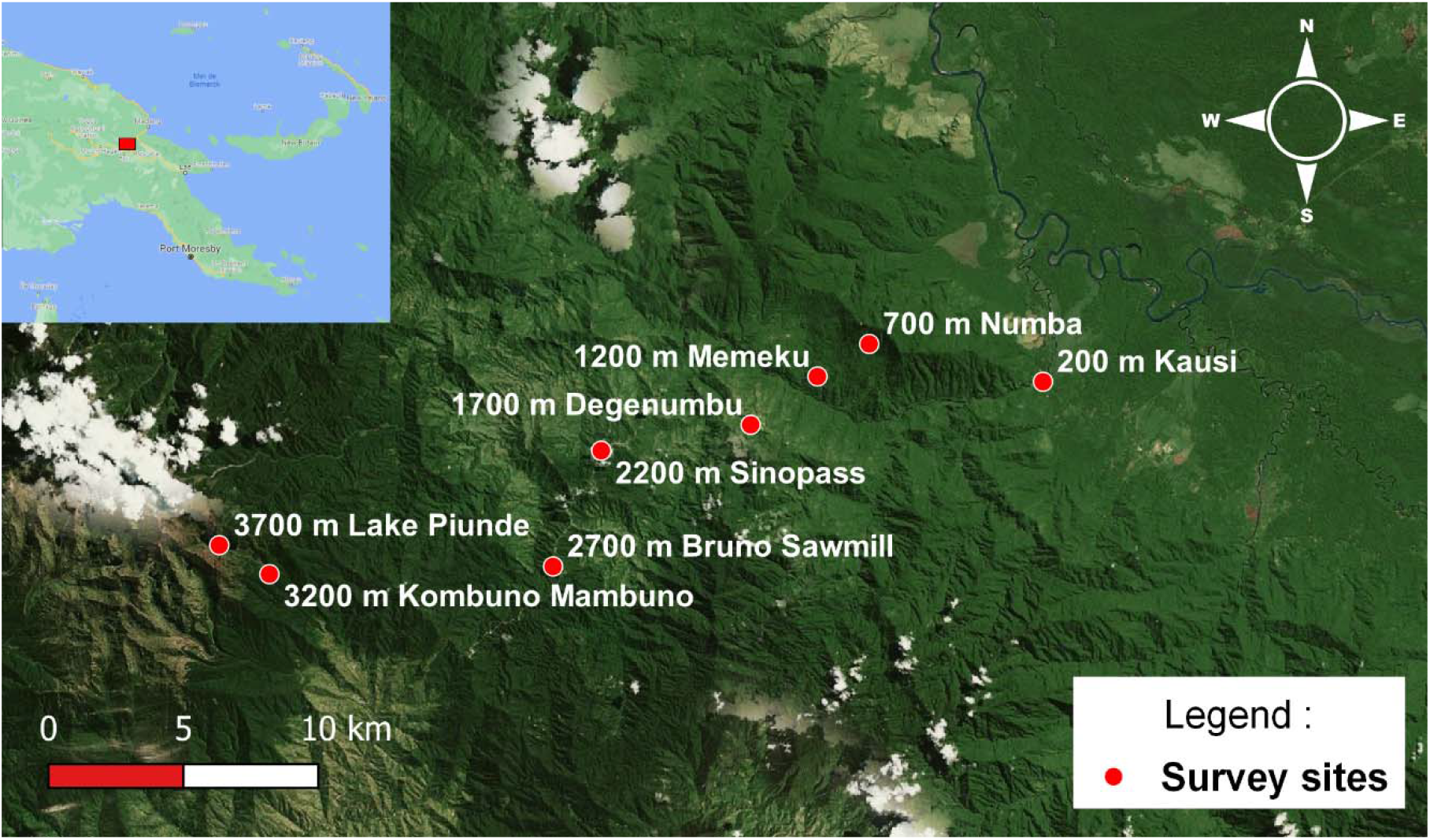
Map of the Mt. Wilhelm elevational gradient (source: Google earth; QGIS 3.12.2) and its location in Papua New Guinea (insert with red square).

### 2.2 Mist-netting and acoustic surveys

The bat communities were surveyed by mist-netting and by acoustic surveys, during two independent expeditions conducted in wet (February – April 2015) and dry seasons (June – July 2015).

We used portable ultrasonic bat call detectors (Wildlife Acoustics EM3+ and a Pettersson Elektronik D240X connected to a Roland R-05 Wave/MP3 recorder) to record the calls of echolocating bat species. We surveyed the bats at five points separated by 200 meters at each elevation, visiting each of these points for 15 minutes daily (Table S2.1). The points were located along the point-count track of 16 points (150 m apart) used in the previous study of bird communities (Sam et al., 2019). Surveys were conducted for four days per site from February to July 2015 after sunset (6 pm) but were only feasible for two days at 3,200 and 3,700 m. Recordings were analysed by opening each WAV file in Adobe Audition version 22.0 and inspecting the spectrograms for bat echolocation pulses. There were three different sampling rates in the data set: 22.05 kHz, 44.1 kHz, and 256 kHz. The characteristic frequency of echolocation pulses was determined after accounting for the sampling rate and was estimated from the power spectrum. Identification of bat species was undertaken in two steps. First, ‘echolocation call types’ were recognised from the recordings and defined based on a standardised naming scheme that has been used in many published and unpublished surveys across Papua New Guinea and Wallacea in recent years (Armstrong et al., 2020, 2015a, b; Armstrong & Aplin, 2011, 2014; Kale et al., 2018) and second, bat species identifications were attributed to each echolocation call type based on information from these and other surveys (annotated species list in Table S2.2 and illustration of call types in Figure S2.2). This two-step approach, along with the provision of illustrated examples of identified call types, provides transparency that allows for future verification of species identifications, and retrospective correction of species names on the basis of updated information. Nomenclature follows the current IUCN Red List profile for each species (https://www.iucnredlist.org/).

We set a total of eight mist-nets (length x height: 15 x 3.5 m) in various habitat types per study site, including understorey ‘flyway’ spaces along human tracks, across creeks, and forest openings. We mist-netted five nights (12 hrs per night) per site in the wet season survey. During the dry season, we revisited elevations from 200 to 2,700 m only, and we operated the mist nets over four nights for five hours daily. Mist-nets were moved to a new spot after every two or three nights. All mist-netted bats were identified to species using field guides by Bonaccorso (1998) and Flannery (1995) as well as (Irwin, 2017; Parnaby, 2009) for *Nyctophilus timoriensis* and *Nyctimene cyclotis* (Figure S2.1). However, morphologically similar *Paranyctimene raptor* and *P. tenax* could have been misidentified in the field; they can occur in sympatry, as previously observed by Bergmans (2001).

### 2.3 Regional data and explanatory variables

The regional data included only bat species described as present in Central Range and Sepik- Ramu Basin in Bonaccorso (1998). We summarized each bat species’ food preferences and elevational ranges described in the book (Table S3.1). The elevational ranges attributed to the bat species came from captures across the whole New Guinea. Nevertheless, it still reflects their tolerance to elevation even though some bat species are not found across the entire PNG area within their elevational range.

We used the log_transformed available land area of elevational belts 200 m wide across the whole New Guinea mainland as a proxy for the land area available for respective study sites (e.g., 100–300 m a.s.l., for the 200 m a.s.l. study site; measured in GIS software ARCGIS 9,3 and ERDAS ER Mapper 6). Indeed, an elevation belt of 200 m seems to perform the best as a proxy of the available land area to explain species richness patterns, according to a recent publication (Sam et al., 2019).

Temperature and humidity were recorded every hour for one year (April 2010–July 2011) using a data logger (Comet R3120) placed in the forest interior of each study site. Mean annual temperature decreased at a constant rate, while mean humidity remained high across the entire transect (83.0%–97.4%) (Table S3.2). Thus, we used only mean daily temperature as a predictor variable.

We used three variables related to habitat (Table S3.2), measured at each point (i.e., 16 points per elevational study site, 128 in total, Sam et al., 2019): (1) Average canopy height (using a laser rangefinder; three measures/point), (2) Shrub density (using a vegetation board (Lilith, 2007; MacArthur & MacArthur, 1961), five measures/point, 1–3 m height), and (3) Canopy openness (5 photos/point analysed with a Gap Light Analyzer; Frazer et al., 2001). These three factors determine the vegetation structure at each site, which is the basis for the organization of bat foraging guilds, defining flight spaces for foraging bats, and controlling the availability of roost sites (Charbonnier et al., 2016; Denzinger & Schnitzler, 2013; López_González et al., 2012).

In addition, we derived several predictors of food availability. Firstly, we used two food variables for frugivorous-nectarivorous bats (Table S3.2), (1) species richness, and (2) abundance of fruiting trees (Villemant et al., 2016). Trees were counted and identified in three random plots of 20 x 20 m at each study site in the dry season of 2013. Further, we obtained the abundance and richness of the fruiting trees targeted by bats at each elevation by using plant genera recognized as having fruit or nectar consumed by pteropodid bats by the database of Aziz et al. (2021).

Moths (Lepidoptera) dominate the diet of most insectivorous species. We used the (1) species richness and (2) abundance of Geometridae, one of the most important moth families (Beck et al., 2017; Vestjens & Hall, 1977) as an indicator of Lepidoptera availability. The specimens were collected using manual light trapping (May–August 2009; October–December 2009, January 2010) at all eight study sites of Mt. Wilhelm transect (Beck et al., 2017; Toko, 2011).

### 2.4 Statistical analysis

We used incidence data (i.e., presence/absence per sampling night) from the acoustic surveys as it was not possible to separate the vocalizations of individual bats. Mist-netting data were also converted to incidence data to facilitate comparison. We recorded the number of nights at a particular elevation when a given species was present. We used sample-based rarefaction to compare species richness by sampling days at each elevation (in EstimateS 9.1; Colwell, 2013) for both methods. We extrapolated the sampling effort by doubling the number of sampling days. Then, we used non-parametric estimators for incidence data (Jackknife1) to compare with the richness observed along the transect using the software EstimateS 9.1 (Colwell, 2013). Abundances (from mist-netting data) and total species richness (from both methods) were also described for each of the three feeding guilds (frugivore-nectarivore, insectivore, frugivore- nectarivore-insectivore).

We produced heatmaps using the Jaccard dissimilarity index to compare bat species composition between sites using the “Vegan” library (Oksanen et al., 2013) in R software (R Core Team, 2020). The first heatmap was run using incidence data from both mist-netting and acoustic surveys. The second one was produced with regional data (i.e., bats from Central Range and Sepik-Ramu basin). We calculated the mean elevation range for all species observed at each elevation in this study and from regional data, then compared local and regional ranges.

Total species richness was used as the dependent variable in Poisson regressions with combinations of five predictor variables (log-transformed): mean daily temperature, canopy height, shrub density, canopy openness, and land area available within 200 m elevational belts (Table S5.1). Similarly, species richness partitioned into feeding guilds was used as the dependent variable with combinations of six variables: mean temperature, land area, fruiting tree richness, and abundances, or moth species richness and abundances. The small size of our dataset did not allow us to test habitat variables in partitioned richness and to include interactions in our models. We used ΔAICc and Akaike weights (w_1_) to interpret regression results and evaluate models and their fits (Anderson & Burnham, 2002). Frugivore and nectarivore species were assigned to one main feeding guild in our models: frugivore-nectarivore. Frugivore-nectarivore-insectivore species (i.e., *Syconycteris australis*) was included into two guilds —the frugivore- nectarivore and insectivore guilds (i.e., its presence was included in two datasets).

## 3. RESULTS

### 3.1 Species richness pattern

By mist-netting, we captured 701 individuals of 12 bat species (Table S2.1). We did not capture any individuals above 2,700 m during the wet season and we were not able to resurvey elevations above 2,700 m during the dry season due to logistical constraints. A total of 11 echolocation call types were recognised from the recordings, each of which can be associated with one or more bat species (Figure S2.2). From these, at least ten bat species in five families were recorded (i.e., confirmed) as being present on the survey (Table S2.1). In total, no less than 22 species were observed in five months along the Mt. Wilhelm transect. This represents about 30 % of the regional species pool according to Bonaccorso (1998). Species richness declined with increasing elevation, regardless of the survey method used or the data sources (Figure 2).

**Figure 2:**
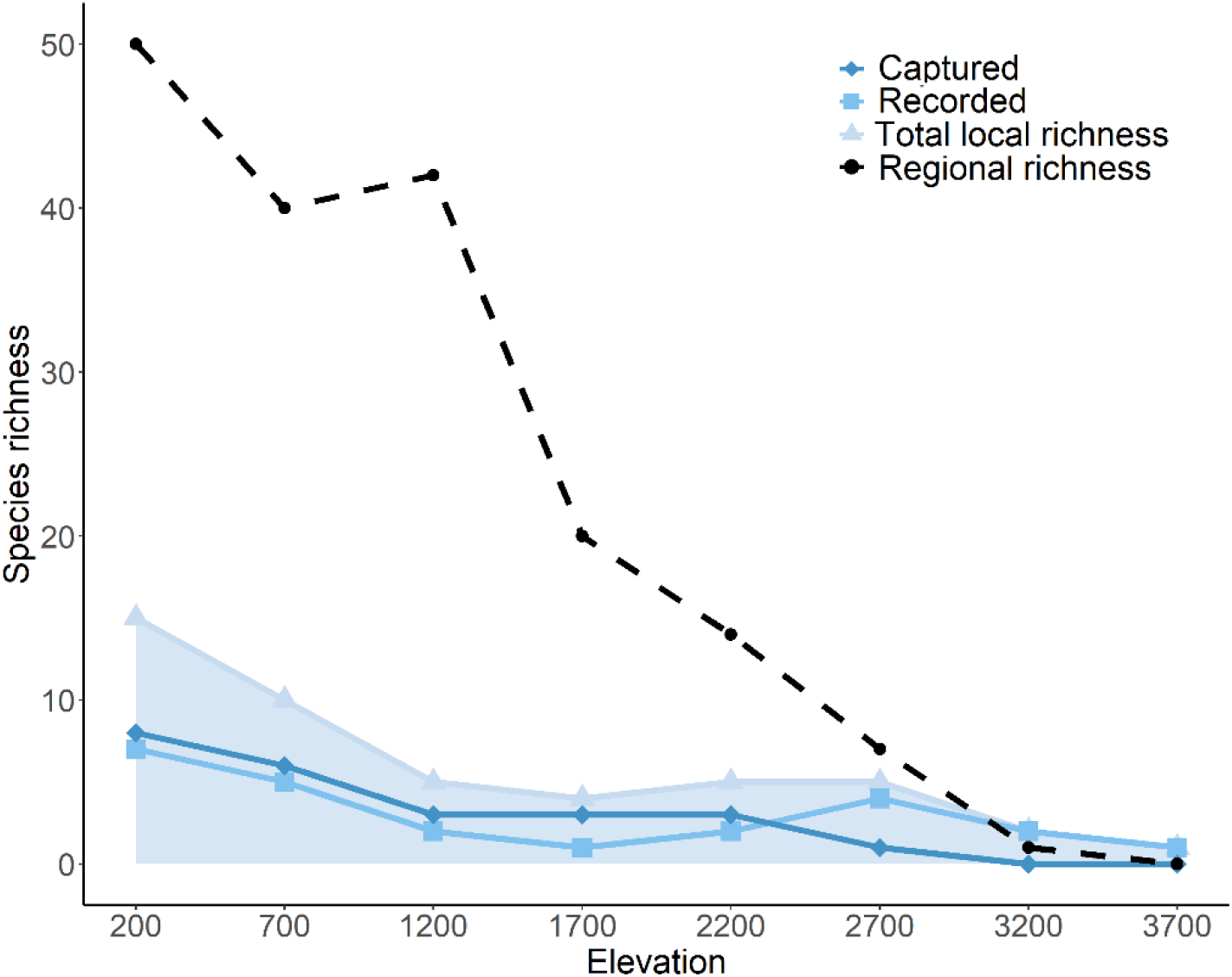
Bat species richness recorded from mist-netting captures, acoustic records, and both methods combined, at eight study sites of Mt. Wilhelm elevational transect in Papua New Guinea. Regional species richness according to Bonaccorso (1998).

The non-parametric estimator “Jackknife 1” matched the species richness trends, but its estimates were higher than the observed curves (Figure 3c, 3d). Specifically, it predicted significantly higher than observed richness at 700 and 2,700 m in audio data and from 200 m to 2,200m in mist-netting data. The species accumulation curves have not reached the plateau during the acoustic surveys at 700 and 2,700 m and by mist-netting at 700 m (Figure 3a, 3b). We noticed a low plateau at 1,700 m in acoustic data and 1,200m in mist-netting data which matched with the decreases observed at these elevations on the species richness trends (Figure 3c, 3d). On the other hand, the rarefaction curves showed that we quickly reached the highest level of species richness at high elevations (3,200–3,700 m) in both survey methods.

**Figure 3:**
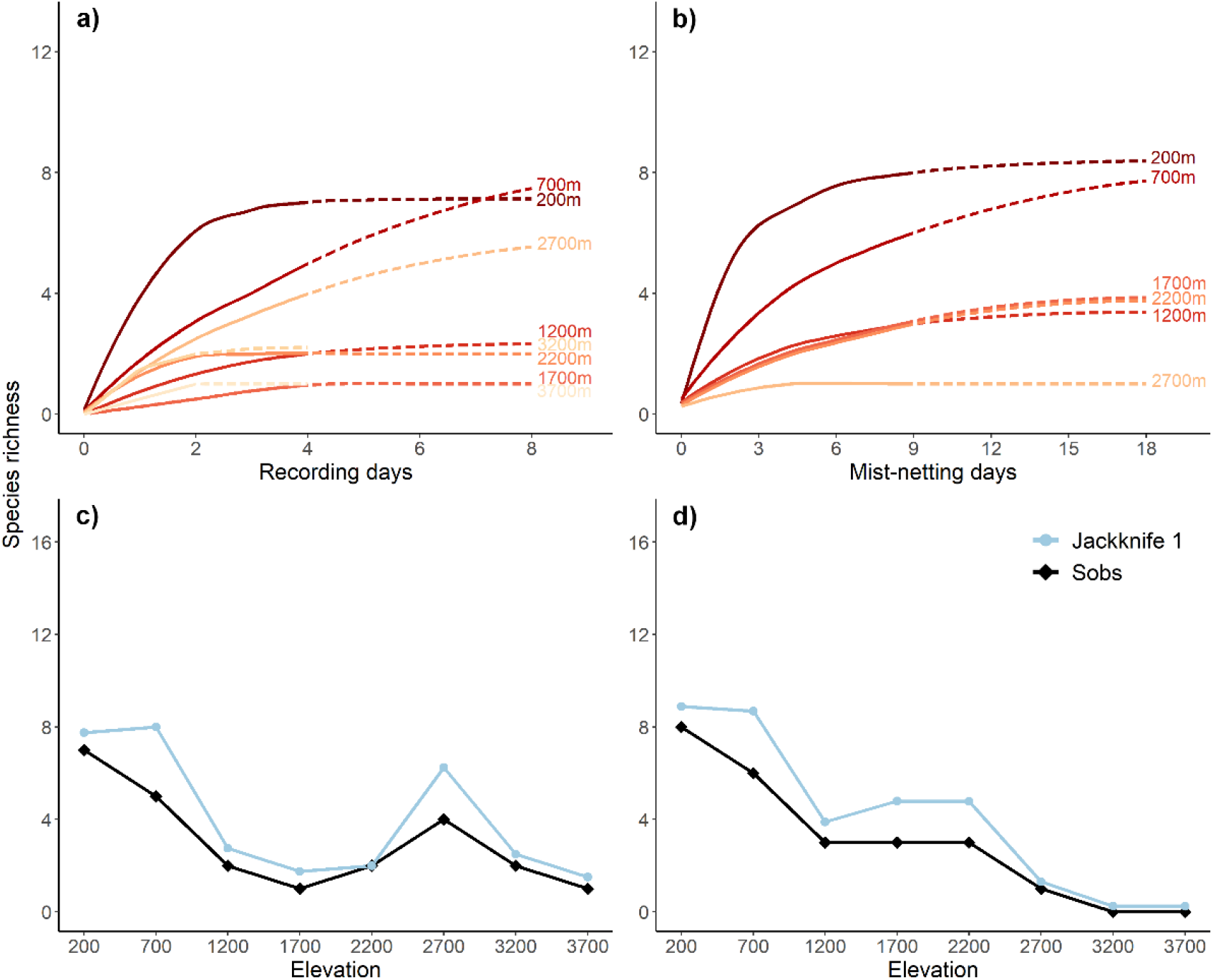
Rarefaction curves of observed and extrapolated (dashed lines) (days x2) number of bat species according to the number of recording (a) or mist-netting days (b) along the elevational transect of Mt. Wilhelm in Papua New Guinea. Observed, estimated (Jacknife1) species richness using audio (c) or mist-nets (d) along elevational transect of the Mt Wilhelm.

The mist-netting data did not reveal a clear pattern of abundances along the elevational transect (Figure S4.1) while the acoustic data could not be used to estimate abundance. Fruit-feeding bats were the most frequently captured (Figure S4.1) and *Syconycteris australis* dominated the samples (Table S2.1). However, in terms of the number of species, the richness of the frugivores-nectarivores and insectivores declined in a similar pattern along the transect (Figure S4.1).

### 3.2 Drivers of diversity

We first modelled total species richness with the abiotic and habitat variables, followed by partitioned species richness (i.e., frugivore-nectarivore, insectivore) with the abiotic and food variables. Selected according to AICc, the model with the available land area as a single variable performed better than any other combination (Table S5.1) when it included all species or only insectivores, while temperature as a single variable performed better in frugivore species (Table 1). Despite strong correlations between canopy openness (R>0.8) and height (R>0.8) with area and temperature (Figure S3.1), habitat (Figure S6.1) and any food variable (Figure S6.2 and S6.3) did not significantly improve the model.

**Table 1:** Akaike’s second□order information criterion (AICc) for single-predictor models of observed bat species richness along the Mt. Wilhelm elevational transect, estimated for all bat observations and the observations partitioned into two feeding guilds. The bold text underlines the model which performed better than any possible combination. Note that only single-predictor models are shown here, while combinations of all the explanatory variables are available in Table S5.1 in Appendix S5.

| *All bats* | $-\log(L)$ | Akaike weight ($w_i$) | $AIC_c$ | $\Delta AIC_c$ |
| --- | --- | --- | --- | --- |
| **Area** | **14.55** | **0.458** | **35.5** | **0.00** |
| Temperature | 15.32 | 0.212 | 37 | 1.54 |
| Canopy height | 16.31 | 0.079 | 39 | 3.52 |
| Shrub density | 18.79 | 0.007 | 44 | 8.49 |
| Canopy openness | 18.85 | 0.006 | 44.1 | 8.61 |
| *Frugivores-nectarivores* | $-\log(L)$ | Akaike weight ($w_i$) | $AIC_c$ | $\Delta AIC_c$ |
| **Temperature** | **9.31** | **0.552** | **25** | **0.00** |
| Area | 9.95 | 0.289 | 26.3 | 1.29 |
| Fruiting tree abundances | 15.65 | 0.001 | 37.7 | 12.67 |
| Fruiting tree richness | 16.54 | 0.000 | 39.5 | 14.46 |
| *Insectivores* | $-\log(L)$ | Akaike weight ($w_i$) | $AIC_c$ | $\Delta AIC_c$ |
| **Area** | **13.99** | **0.482** | **34.4** | **0.00** |
| Temperature | 14.42 | 0.312 | 35.3 | 0.87 |
| Moth abundances | 17.67 | 0.012 | 41.7 | 7.36 |
| Moth richness | 17.74 | 0.011 | 41.9 | 7.5 |

### 3.3 Assemblages of bat communities

The heatmap based on the Mt. Wilhelm data (Figure 4a) showed a rapid turnover of communities along the elevation transect. We found high similarity between pairs of communities at the extreme ends of the transect, between 200 and 700 m and between 3,200 and 3,700 m but also at mid-elevation between 1,200 and 1,700 m. The latter pair of sites shared the only species (i.e., *Miniopterus* australis) found at 3,700 m with a total of two species at 3,200 m. However, using regional data (Figure 4b), the heatmap revealed an increasing turnover of species with increasing elevation. Communities also appeared similar between the closest sites (e.g., 200–700 m; 700–1,200 m) except for the highlands. The elevational distribution of bats ends at 3,200 m in regional data (Table S3.1) so that the highest elevation (3,700 m) could not share any species with other sites.

**Figure 4:**
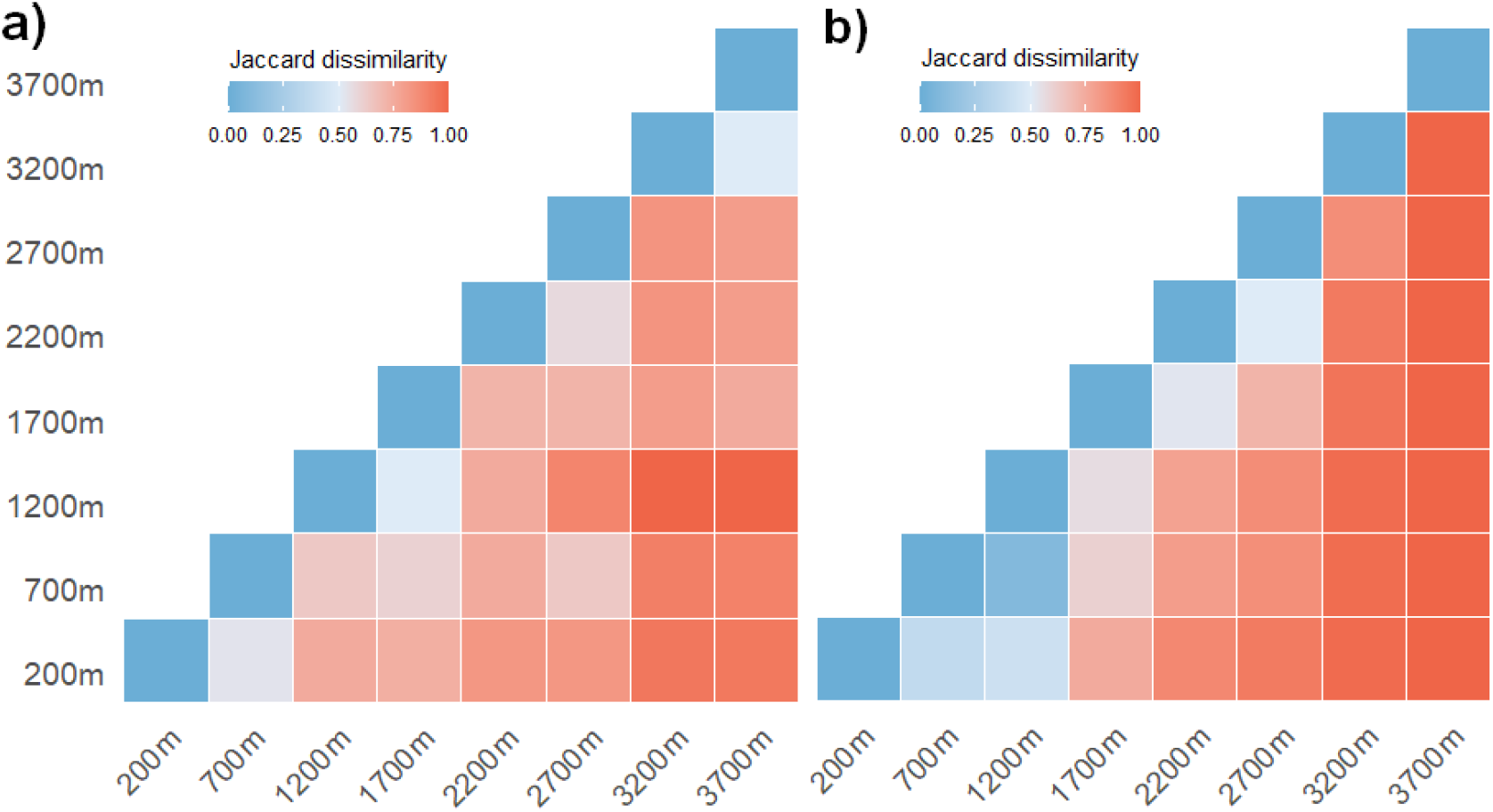
Heatmap of Jaccard dissimilarity index of local (mist-netted and acoustic data combined) bat communities found at Mt. Wilhelm (a) and potential bat communities based on the regional distribution of bats in Papua New Guinea (b).

In the regional data, the mean elevation range length of species is increasing with increasing elevation, meaning that species occurring at one elevation tend to be also found at the lowest ones (Figure 5). The same pattern was observed in our community samples except at 3,200 m where it decreased slightly. It means that one of the species recorded by us (i.e., *Miniopterus tristis/Pipistrellus collinus*) at 3,200 m was not detected in the lowlands, despite occurring there in the regional data.

**Figure 5:**
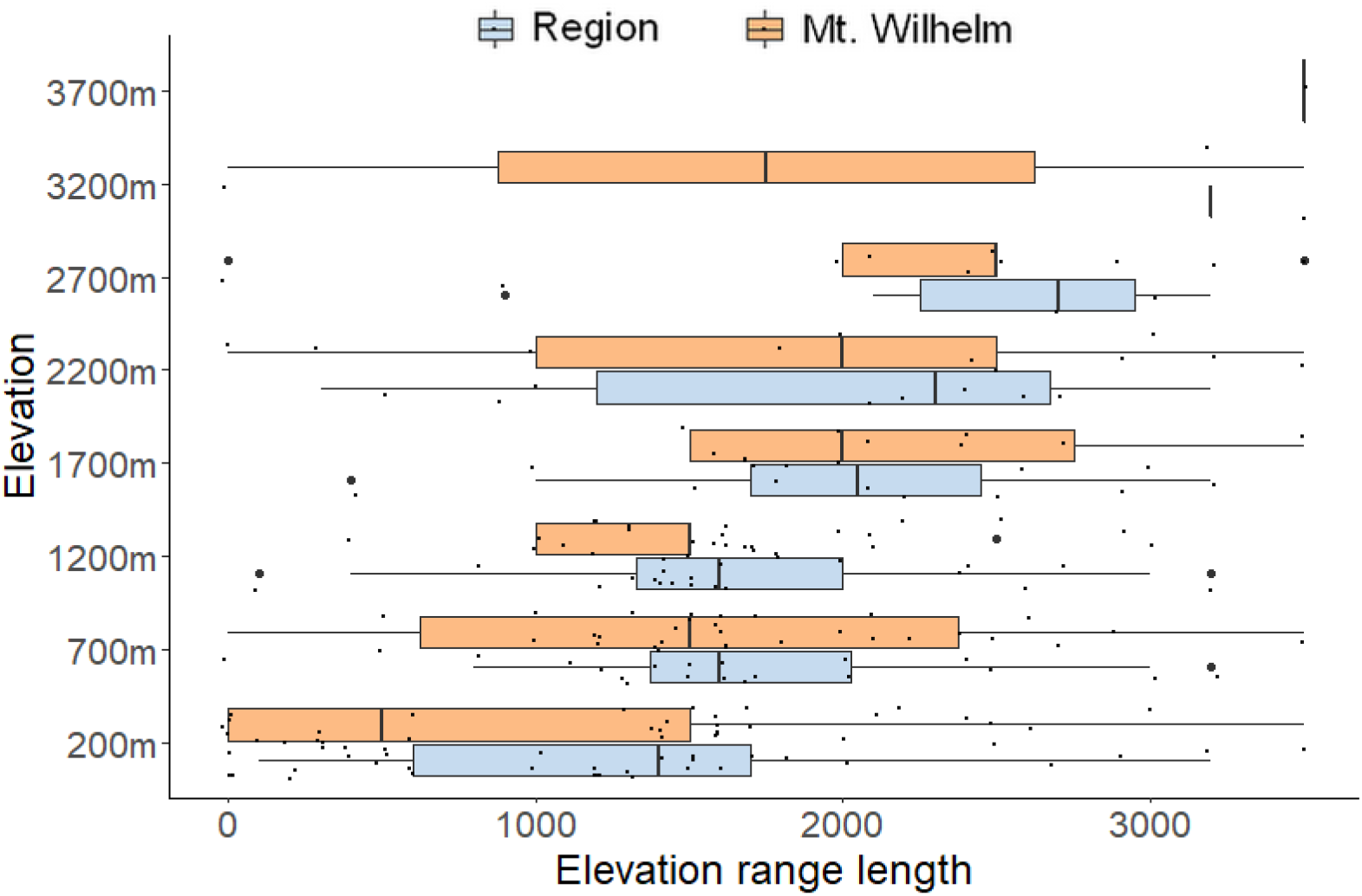
The elevation range for all bat species occurring at each surveyed elevation calculated from Mt. Wilhelm and regional data. Note that only one species is recorded at 3200 m and none at 3700m in the regional data, while we recorded two and one bat species at 3200 and 3700 m respectively (Amick et al., 2021).

Our results confirmed the upper elevation reported in the literature for most of the species at Mt. Wilhelm, with the exception of six species: *Hipposideros wollastoni, H. calcaratus, Dobsonia minor, Nyctimene draconilla, Paranyctimene raptor, Miniopterus australis* [=*Miniopterus* sp. 2 ‘small’]), that were observed at higher elevations (Figure S7.1).

## 4. DISCUSSION

Our study was the first detailed bat survey along a forest transect in PNG and revealed that Mt. Wilhelm hosts at least one-third of the bat species richness expected in that region (Bonaccorso, 1998), representing thus an important diversity hotspot. Bat species richness decreased with increasing elevation and the observed pattern was mainly driven by available land area or mean daily temperature. Species turnover was continuous, with most of the bat species occurring in lowlands and only a limited number reaching the higher elevations. However, while this pattern was predictable based on the regional data, our observations at Mt. Wilhelm showed that some bats were detected only at high elevations and seemed to be missing in the lower parts of their typical ranges. This pattern might be real or affected by sampling effort issues. Extensive studies are required to clarify this pattern and understand bat community structures in PNG.

### 4.1 Species richness pattern

PNG is home to a very high number of bat species (95 species) and, with 36 species, boasts the second-highest diversity of Old World fruit bats (i.e., family Pteropodidae) in the world after Indonesia (Aziz et al., 2021). However, it is also one of the most data deficient and poorly-understood countries in terms of how the bat communities are structured with respect to their ecological roles (Aziz et al., 2021). Mt. Wilhelm represents a high diversity of bats with at least 22 species out of the 62 expected in that region. Our data indicate that more species would be revealed by a more intensive survey.

The decreasing species richness with increasing elevation followed the main trend found in tropical mountains documented by McCain (2007). As expected, we confirmed differences between results provided by the two sampling methods employed (Kuenzi & Morrison, 1998; Larsen et al., 2007) whereby most of the species captured in the mist nets were frugivores and nectarivores. Insectivorous bats possess specialized anatomical adaptations that enable echolocation and thus, some species are able to detect and avoid the mist-nets (Francis, 1989; Larsen et al., 2007). This likely explains why only two echolocating bat species were captured in our study, in addition to being detected acoustically (Table S2.1 in Appendix S2). The rarefaction curves showed that acoustic surveys would have to be longer than eight days and coupled with all-night recordings from autonomous recorders in order to yield accurate numbers of species at 700 and 2,700 m. Furthermore, a recent study in the Neotropics revealed that the main centre of activity in bat species is in rainforest canopies (Marques et al., 2016), which has already been observed for birds in PNG (Chmel et al., 2016). However, since we did not survey bats at the canopy level due to logistical challenges, we potentially missed bats flying above the treetops, perhaps because our equipment could not detect the attenuated echolocation signals, or they were simply less likely to reach our understorey mist nets (Kalko & Handley, 2001; Marques et al., 2016). Our result underscored the tendency of insectivorous species to avoid mist-nets and the necessity to employ both acoustic and capture methods in the forest canopy and understorey.

In terms of abundance, we captured more frugivore-nectarivore individuals in the lowlands (i.e, 200–700 m) but we did not see significant differences in the capture rate from 1,200 to 2,200 m. This contrasts with studies in South American forests (Carvalho et al., 2019) where frugivores declined greatly above 1000 m. At 3,200 to 3,700 m, the habitats along rivers were mostly open (Figure S1.1 in Appendix S1), which made the captures more difficult. That is perhaps why we did not catch any bats above 2,700 m. However, we have no independent way of assessing catchability in the frugivorous species, in contrast with the insectivorous species for which we can detect them acoustically even though they are not easily captured. Therefore, we were unable to resolve whether frugivore-nectarivore are simply missing above 2,700 m due to environmental filters (e.g., fruit production, temperature) or whether it is the result of apparent mist-netting limitations.

### 4.2 Drivers of diversity

Despite using a range of factors including some rarely considered for bats (i.e., food availability, habitat), our analysis revealed that available land area and mean daily temperature have the strongest effect on bat species richness along the elevational transect. The area available in the 200 m elevational belt around the study site was the best predictor in the models including all the species or insectivores only, followed closely by temperature in explanatory power. It followed the long-known and very robust pattern of the species-area relationship (May, 1975; Rosenzweig, 1995), where the number of species increases with the size of the area.

Temperature was the best predictor of frugivorous species. As such, our data mirror those from other studies conducted in tropical mountains where temperature was also reported to be a strong correlate with bat diversity (McCain, 2007b). Temperature could affect distributions through direct (e.g., physiology) or indirect effects (e.g., habitat, food resources) in different ways between feeding guilds. Mean daily temperatures were 9.9 and 7.9 °C at 3,200 and 3,700 m respectively, which is below the temperature tolerance of most bat species (Appel et al., 2019; Turbill, 2008) and may explain why we did not capture any bats at these sites, and only recorded two species above 2,700 m. Besides, temperature could influence bat species richness indirectly through vegetation and food resources (Charbonnier et al., 2016; Moura et al., 2016). Indeed, vegetation structure could be important for bats, indirectly related to food resources, but also for roosting sites (Capaverde et al., 2018; Kunz, 1982; Perry et al., 2007). Roosting opportunities depend on the number of trees of appropriate size and whether or not they contain cavities or other structures appropriate for bats. Based on published data, 67 % of the species detected in Mt. Wilhelm potentially use foliage or tree hollows (Table S8.1 in Appendix S8). In addition, insect abundance, and fruit and nectar production are all predicted to be low at high elevations (Loiselle & Blake, 1991; Terborgh, 1977). Studies suggest that the reduction in productivity with elevation (McCain & Grytnes, 2010) has a more substantial impact on fruit resources (e.g., figs) than on the other types of resources used by bats (e.g., insects) (Presley et al., 2012; Segar et al., 2017). The distribution of moths from Mt. Wilhelm shows a mid-peak pattern (Beck et al., 2017; Toko, 2011), which does not seem to be followed by insectivorous bats. This is in contrast with patterns in abundances of insectivorous birds of Mt. Wilhelm which peaked in abundance at mid-elevations, where habitat complexity was the greatest and arthropod abundances were still relatively high (Sam et al., 2020).

The inclusion of habitat features, fruiting tree and moth species richness, and abundances in the model did not change the relative level of influence of temperature and area on species richness. However, available area, as well as temperature, decrease linearly with increasing elevation while bat diversity does not. The steepest drop-off in numbers was between 200 m and 1,200 m, and declining at a slower rate thereafter. It is thus likely that there are also other factors, that we did not consider (e.g., seasonal food availability), modifying the response by bats.

### 4.3 Assemblages of bat communities

According to regional data, species turnover kept increasing from similar communities in the lowest study sites (200–700 m) to more dissimilar ones in the highest sites (2,200–3,200 m). Nevertheless, the communities of bats found at the highest sites were just a subset of bats from the lowlands, suggesting that the vast majority of the bat species found in this region are not restricted to mountainous areas. However, potential identification issues and unresolved taxonomy might affect the understanding of species elevational distributions, especially in this long-standing regional dataset (Bonaccorso, 1998).

Our study is the first bat survey conducted at the highest peak of PNG. We detected bats at the 3700 m elevation band—never before recorded in PNG (Amick et al., 2021)—and observed wider bat ranges than the ones described previously for the region (Bonaccorso, 1998). These range extensions are most likely due to low sampling effort for bats at high elevations in PNG in the past, rather than any recent range expansions. In Mt. Wilhelm surveys, we found bat species at high elevations that were missing from the lowlands (M. tristis/P. collinus, H. wollastoni, O. secundus), which have been encountered more commonly at mid-high elevations. Globally, previous studies showed that the majority of bats found in highlands are primarily lowland species that occasionally commute to higher elevations when conditions become favourable (Alberdi et al., 2015; Michaelsen, 2010) or use these environments as commuting routes. Indeed, as previously mentioned, most bats are limited by direct and/or indirect effects of temperature, and life at high elevations could present an energetic challenge, especially for pregnant and lactating females (Kunz et al., 1995). In our sampling, low canopy openness in the lowlands could affect our ability to detect echolocation signals from bats flying over the canopy. Moreover, changes in call structure at high elevations have already been observed for one bat species (i.e., *Tadarida brasiliensis*) in a previous study (Gillam et al., 2009). Nevertheless, under a scenario of climate change, these bat species (i.e., M. tristis/P. collinus, H. wollastoni, O. secundus) may have been recently constrained to high elevations because of spatial elements and/or their ecology and/or physiology. However, we are unable to resolve whether we likely missed these species in the lowlands because of the use of methods that have a greater bias towards the detection of species below the canopy or whether these bat species spanning only in high elevations are the consequences of acclimations to climate change. Extensive surveys with a greater level of effort might help to answer this question.

## 5. CONCLUSION

Mt. Wilhelm provides habitat for a globally significant bat fauna whose species richness follows the typical decreasing pattern with elevation found in other tropical mountains. Available land area and mean daily temperature explained the vast majority of this pattern, however, we suspect that additional factors (i.e., seasonal food availability) could improve the models. Bat communities also varied gradually along the elevational transect, as we describe for the first time for PNG. The fact that six species in this study were recorded above their typical elevational range (Table S2.1 in Appendix S2), including some detected above the previously described maximal distribution for PNG, might be the result of climate change or basic survey issues and lack of good quality historical data. A greater level of effort would shed light on this, which is potentially important, as the bats of PNG remain largely understudied. This study highlights how the results of basic inventory surveys that employ a comprehensive, multi-method effort for bat sampling can be used to address ecological questions that might help with impact assessments.

## ACKNOWLEDGMENTS

We thank the landowners of Mt Wilhelm elevational transect and villagers from Bruno Sawmill, Sinopass, Degenumbu, Memeku, Numba, and Kausi for assistance and access to the sites. We also thank P. Toko who shared his database on moth species along Mt Wilhelm elevational transect, and L. Sam and R. Hazel who shared data on the richness of fruiting trees. Binatang Research Centre helped with logistical support and research permits. We acknowledge ERC StG BABE 805189 and GAJU n.04-048/2019/P for financial support. Bat sampling was conducted under research permit 11800056119.

## AUTHORS CONTRIBUTIONS

PKA conducted the fieldwork, recorded bat calls, and mist-netted the bats, KNA performed bat call identifications, ES performed data analyses, extracted data from literature and wrote the first draft of the manuscript, KS designed and funded the study and helped with the analyses, PKA, KNA, KS, and VN contributed significantly to revisions.

## DATA AVAILABILITY STATEMENT

The data used for analysis are available in Table S2.1 in Appendix S2 and Tables S3.1, S3.2 in Appendix S3.

## Notes

### Competing Interest Statement

The authors have declared no competing interest.

