## Supporting information for "Species richness and community structure of bats along a forest elevational transect in Papua New Guinea"

**Appendix S1**

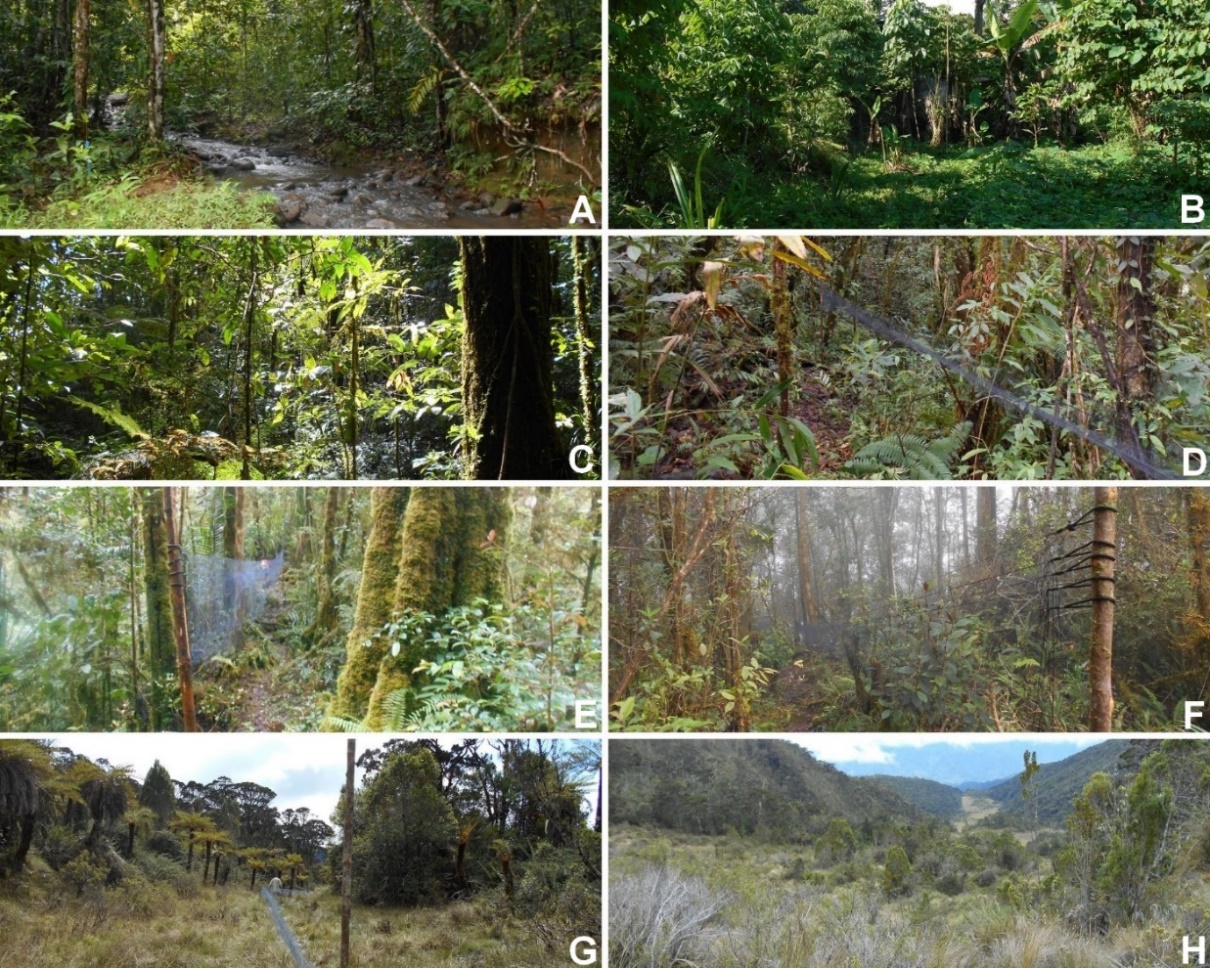

Fig. S1.1: Pictures of the different habitats along the Mt. Wilhelm elevational gradient A : Kausi (200 m) ; B : Numba (700 m) ; C : Memeku (1200 m) ; D : Degenumbu (1700 m); E : Sinopass (2200 m); F : Bruno Sawmill (2700m); G : Kombuno Mambuno (3200 m); H : Lake Piunde (3700 m).

**Appendix S2**

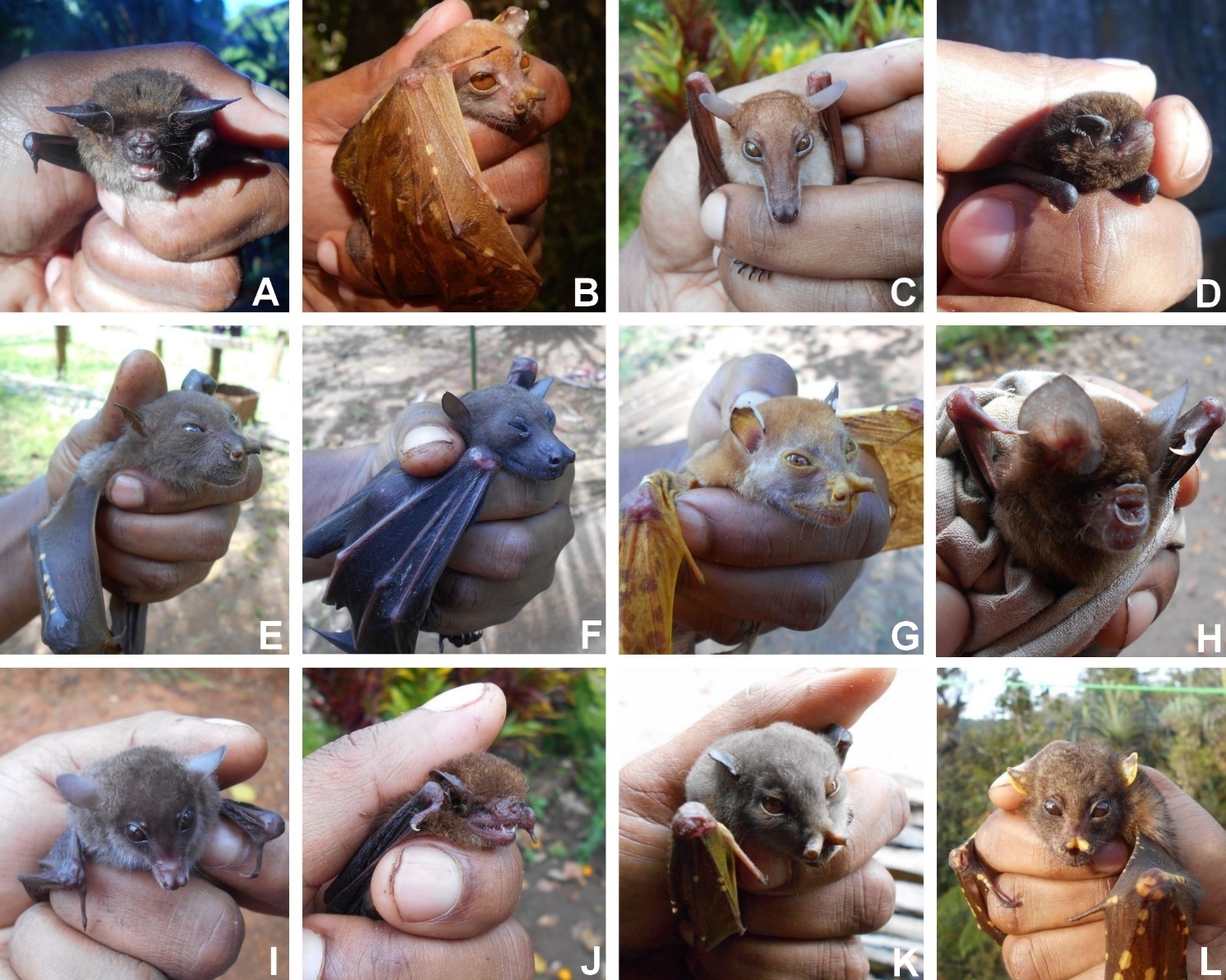

Fig. S2.1: Pictures of the bat species captured using mist nets (source: P. Amick). A : Nyctophilus shirleyae^1^ ; B : Nyctimene albiventer ; C : Macroglossus minimus ; D : Miniopterus australis; E : Nyctimene aello; F : Dobsonia minor ; G : Nyctimene draconilla ; H : Hipposideros calcaratus I : Syconycteris australis ; J : Nyctophilus microtis ; K : Paranyctimene raptor; L : Nyctimene certans^2^

^1^Previously *Nyctophilus timoriensis* (Parnaby 2009)

^2^Previously *Nyctimene cyclotis* (Irwin 2017)

Table S2.1: Bat species captured (MN in Call type column) and acoustically recorded along the Mt. Wilhelm gradient. Grey zones underline bat species captured and recorded as well. Numbers indicate the number of individuals captured. Blue zones underline the bat species elevational ranges previously described in Bonaccorso (1998). Note that call types not attributed to a specific species or more than one species have no blue zones. Feeding guilds come from Bonaccorso (1998). See Fig. S2.2 for illustration of echolocation call types.

| **Species** | **Call type** | **200** | **700** | **1,200** | **1,700** | **2,200** | **2,700** | **3,200** | **3,700** | **Food** |
| --- | --- | --- | --- | --- | --- | --- | --- | --- | --- | --- |
| **HIPPOSIDERIDAE** |  |  |  |  |  |  |  |  |  |  |
| *Hipposideros wollastoni* | *84 mCF* |  |  |  |  | X |  |  |  | Insects |
| *Hipposideros cervinus* | *140 sCF* | X |  |  |  |  |  |  |  | Insects |
| *Hipposideros calcaratus* | *MN* |  | 1 |  |  |  |  |  |  | Insects |
| **EMBALLONURIDAE** |  |  |  |  |  |  |  |  |  |  |
| *Emballonura beccarii /*  *Mosia nigrescens* | *62 i.fFM.d* | X | X | X | X |  |  |  |  | Insects |
| *Emballonura beccarii /*  *Mosia nigrescens* | *72 i.fFM.d* | X |  |  |  |  |  |  |  | Insects |
| **VESPERTILIONIDAE** |  |  |  |  |  |  |  |  |  |  |
| *Pipistrellus papuanus* | *48 st.cFM* | X |  |  |  |  |  |  |  | Insects |
| *Nyctophilus microtis* | *45-50 bFM + MN* | X | 1 | X |  |  |  |  |  | Insects |
| *Nyctophilus shirleyae* | *MN* | 1 |  |  |  |  |  |  |  | Insects |
| **MINIOPTERIDAE** |  |  |  |  |  |  |  |  |  |  |
| *Miniopterus tristis /*  *Pipistrellus collinus* | *38 st.cFM* |  |  |  |  |  |  | X |  | Insects |
| *Miniopterus* sp. 1 'medium' | *43 st.cFM* | X | X |  |  | X | X |  |  | Insects |
| *Miniopterus australis* [=*Miniopterus* sp. 2 ‘small’] | *MN + 54 st.cFM* | X | X |  | 1 | 1 | X | X | X | Insects |
| **MOLOSSIDAE** |  |  |  |  |  |  |  |  |  |  |
| *Austronomus kuboriensis* | *13 cFM* |  | X |  |  |  | X |  |  | Insects |
| *Otomops secundus?* | *18 cFM* |  |  |  |  |  | X |  |  | Insects |
| **PTEROPODIDAE** |  |  |  |  |  |  |  |  |  |  |
| *Syconycteris australis* | *MN* | 79 | 211 | 89 | 136 | 78 | 19 |  |  | Fruit/ Nectar/ Insects |
| *Macroglossus minimus* | *MN* | 2 | 5 |  |  |  |  |  |  | Nectar |
| *Paranyctimene raptor* | *MN* | 22 | 5 | 6 | 1 |  |  |  |  | Fruit |
| *Nyctimene albiventer* | *MN* | 27 |  |  |  |  |  |  |  | Fruit |
| *Nyctimene draconilla* | *MN* | 3 |  |  |  |  |  |  |  | Fruit |
| *Nyctimene aello* | *MN* | 4 |  |  |  |  |  |  |  | Fruit |
| *Nyctimene certans* | *MN* |  |  | 1 |  | 1 |  |  |  | Fruit |
| *Dobsonia minor* | *MN* | 4 | 1 |  |  |  |  |  |  | Fruit |

Table S2.2 Annotated species list provided to justify identifications and range extensions, with references additional to the main list.

**EMBALLONURIDAE Sheath-tailed bats**

**Beccari’s sheath-tailed bat *Emballonura beccarii meeki***

and/or

**Lesser sheath-tailed bat *Mosia nigrescens papuanus***

Echolocation call types: *62 i.fFM.d* and *72 i.fFM.d*

The echolocation call types *62 i.fFM.d* and *72 i.fFM.d* are attributable to *E. beccarii* and *M. nigrescens*, but associating each call type with each species is currently not possible. Both species have a similar elevational range, and a geographical range that includes, or has the potential to include, the study area (Bonaccorso 1998; Armstrong 2021a,b). In addition, these two species have around the same body size (Bonaccorso 1998) and therefore an expected overlap in the range of the characteristic frequency of the strongest, second harmonic of the call. Association of these two call types with a capture is the best way to provide an unambiguous identification. In the analyses, the echolocation call types *62 i.fFM.d* and *72 i.fFM.d* are considered as two species.

**HIPPOSIDERIDAE Leaf-nosed bats**

**Fawn leaf-nosed bat *Hipposideros cervinus***

Echolocation call type: *140 sCF*

This echolocation call type is typical of *H. cervinus* (Armstrong et al. 2020). No faint fundamental frequencies were evident in the examples recorded.

**Wollaston’s leaf-nosed bat *Hipposideros wollastoni***

Echolocation call type: *84 mCF*

This call type was identified to *H. wollastoni* based on the similarity of the characteristic frequency and the relatively long pulse duration to reference call examples associated with captures from elsewhere (K.N. Armstrong unpublished data). This record extends the geographic range of the species as it is represented by the IUCN Red List (Armstrong and Aplin 2021). *H. wollastoni* is one of few hipposiderids that can be encountered commonly at mid-elevations (e.g., Bonaccorso 1998: up to 2,000 m asl; Armstrong and Aplin 2011: 1,600 m asl; Armstrong et al. 2015a: up to 1,900 m asl, probably misidentified therein as *H. muscinus*, incorrect upper elevational limit cited from this study by Armstrong and Aplin 2021; Armstrong et al. 2020: 1,400 m asl). The record from the present study extends the known elevation range of the species (Bonaccorso 1998) by around 200 m.

**VESPERTILIONIDAE**

**Papuan Pipistrelle *Pipistrellus papuanus***

Echolocation call type: *48 st.cFM*

Distinguishing the echolocation calls of *Pipistrellus* from those produced by species of *Miniopterus* is often quite difficult, but the increasing frequency of the terminal portion of pulses can be a feature that allows attribution to *Pipistrellus*. In this case, given distribution records (Bonaccorso 1998), *P. papuanus* is the most like source, but the New Guinea Pipistrelle *P. angulatus* is also a possibility, as is the unidentified bent-winged bat *Miniopterus* sp. 1 'medium'. Capture is required to provide an unambiguous identification of the source of this call.

**Papuan Long-eared bat *Nyctophilus microtis***

Echolocation call type: *45-50 bFM*

The most likely source of this call type is a species of long-eared bat, and *N. microtis* is the most commonly encountered species at relatively low elevations (Bonaccorso 1998; K.N. Armstrong unpublished data) where it was recorded on the present survey. Another possibility are clutter calls of a species of *Pipistrellus* or *Miniopterus*, which can be difficult to distinguish from calls of *Nyctophilus* spp. if call quality is relatively low. Capture is required to provide an unambiguous identification of the source of this call.

**MINIOPTERIDAE Bent-winged bats**

The entire Indo-Australasian radiation of Miniopteridae was recently revised by Wiantoro (2020), though the names applicable to species in Papua New Guinea have yet to be published formally. The present survey recorded three echolocation call types that can be attributed to a species of *Miniopterus*. Indo-Australasian *Miniopterus* can be categorised into three groups based on overall body size, and representatives of two or more size groups are typically found together throughout the Indo-Australasian region. Until formal publication of names for Papua New Guinean *Miniopterus*, and until follow up capture and DNA barcoding for identification can be undertaken in the study area, reference to the echolocation call types encountered there can be considered more appropriate than species names. Applicable names that might have been attributed in the past are discussed for each echolocation call type.

**Greater Melanesian Bent-winged Bat *Miniopterus tristis grandis***

or

**Mountain Pipistrelle *Pipistrellus collinus* (Vespertilionidae)**

Echolocation call type: *38 st.cFM*

This echolocation call type has two possibilities for its source. One is the larger-bodied bent-winged bat species *Miniopterus tristis grandis* that has been recorded as high as 2,700 m asl previously based on capture and genetic evidence (Armstrong et al. 2020). Alternatively, it could be attributable to *Pipistrellus collinus* given previous records of the species as high as 2,800 m asl (Bonaccorso 1998). Reference calls of *P. collinus* are unavailable. Trapping is required to resolve the identification.

**Unidentified bent-winged bat *Miniopterus* sp. 1 'medium'**

Echolocation call type: *45 st.cFM*

This species would have been referred to as either *Miniopterus macrocneme*, *M. medius* or *M. schreibersii* in the past, which have a geographic range that includes the study area (Bonaccorso 1998; Armstrong et al. 2021a,b). The name *M. schreibersii* is no longer used for Indo-Australasian *Miniopterus* (Tian et al. 2004), with the name *M. orianae* used commonly for the taxon on the Australian continent that is also found in New Guinea (e.g. Churchill 2008; Jackson and Groves 2015). The ‘moderate’ characteristic frequency of the echolocation calls is typical of this body size type on the Papua New Guinea mainland, being at least 5 kHz higher on average than larger-bodied species.

**Unidentified bent-winged bat *Miniopterus* sp. 2 'small'**

Echolocation call type: *54 st.cFM*

This echolocation call type is known to be from a small-bodied bent-winged bat species, which is currently referred to as *Miniopterus australis* in Papua New Guinea (Armstrong et al. 2021c). It is encountered commonly, including at higher elevations (Armstrong and Aplin 2011: 2,900 m asl; Armstrong et al. 2015a: up to 1,900 m asl, possibly misidentified as *Pipistrellus collinus* therein; Armstrong et al. 2020: 2,700 m asl).

**MOLOSSIDAE Free-tailed bats**

**Mantled Free-tailed Bat *Otomops secundus***

Echolocation call type: *18 cFM*

The species name given here is a suggested attribution of this echolocation call type given that there is no reference call information from either of the two *Otomops* species in Papua New Guinea. The harmonic profile of the call is typical of a molossid, and the characteristic frequency is too high for *Austronomus kuboriensis*. Further, the high elevation of the record precludes its attribution to *Chaerephon jobensis*, a species of *Ozimops* or *Otomops papuensis* (based on elevation ranges in Bonaccorso 1998). Other records of calls from mid-elevations attributed to *O. secundus* are from unpublished biodiversity surveys (Armstrong 2021c); and some call types suggested to be from *O. secundus* are of slightly higher frequencies than were recorded in the present study (e.g., Armstrong et al. 2015a: 25 kHz; Armstrong et al. 2020: 30 kHz). The nearest confirmed geographic record of *O. secundus* is Tapu, on the Upper Ramu River Plateau, Madang Province (Bonaccorso 1998).

**New Guinea Free-tailed bat *Austronomus kuboriensis***

Echolocation call type: *13 cFM*

*Austronomus kuboriensis* is the only species of bat in Papua New Guinea that emits an echolocation call that has a characteristic frequency of the fundamental as low as 13 kHz. This species has a recorded in the elevational range between 1,900 and 2,800 m asl (Bonaccorso 1998). The single pulse detected in the present study at 700 m could be incorrect, since it is common to see low frequency signals not attributable to bats resembling echolocation calls in bat detector recordings (K.N. Armstrong pers. obs.), but the quality and pulse characteristics certainly provide reasonable evidence. This elevational record would need to be confirmed with a sequence of pulses.

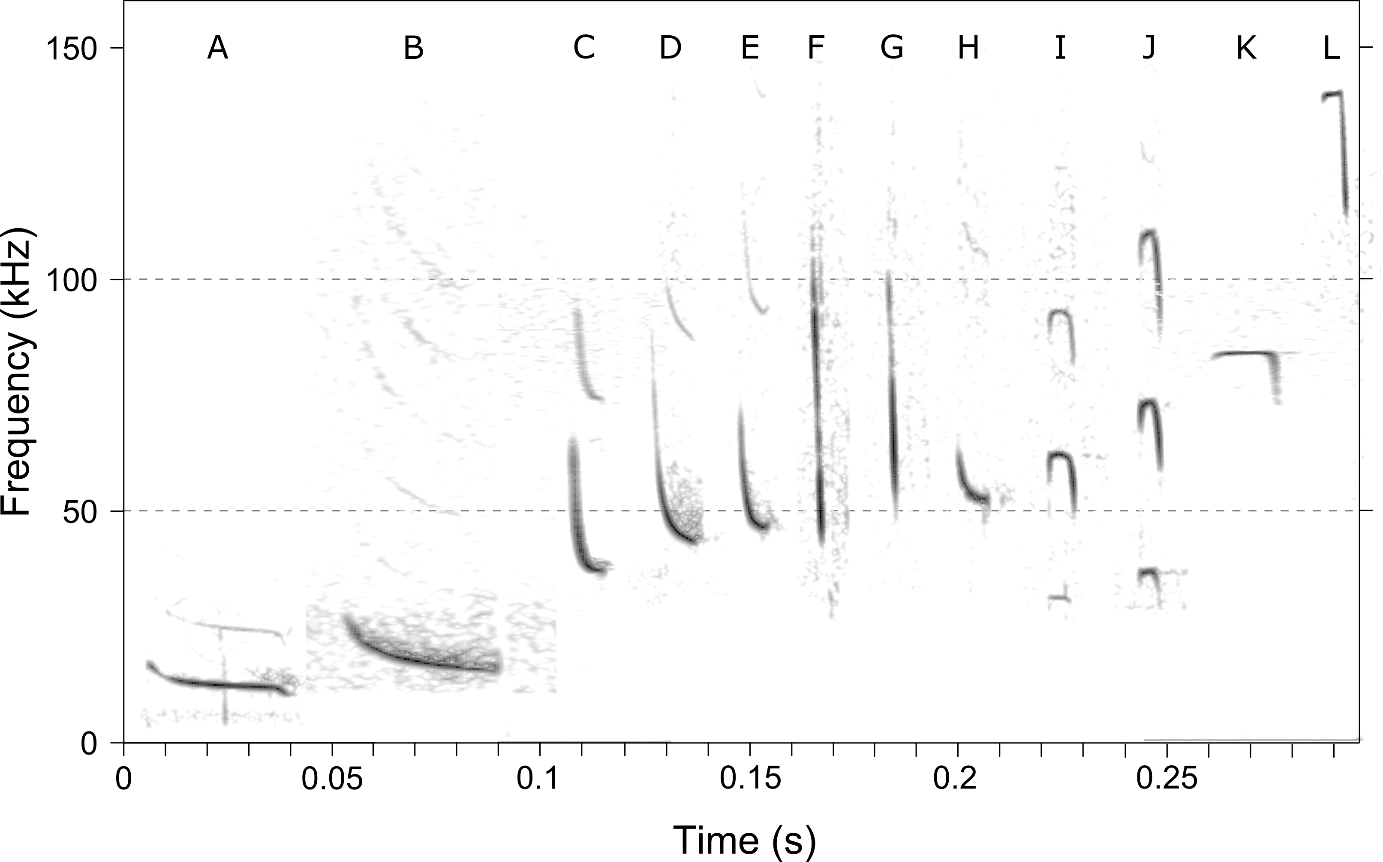

Fig. S2.2. Representative echolocation pulses of the call types recognised and species identified (**A**: *13 cFM Austronomus kuboriensis*; **B**: *18 cFM Otomops secundus*?; **C**: *38 st.cFM Miniopterus tristis grandis / Pipistrellus collinus*; **D**: *43 st.cFM Miniopterus* sp. 1 'medium'; **E**: *48 st.cFM Pipistrellus* *papuanus*.; **F,G**: *45-50 bFM Nyctophilus microtis*; **H**: *54 st.cFM Miniopterus* sp. 2 'small'; **I**: *62 i.fFM.d Emballonura beccarii / Mosia nigrescens*; **J**: *72 i.fFM.d Emballonura beccarii / Mosia nigrescens*; **K**: *84 mCF Hipposideros wollastoni*; **L**: *140 sCF Hipposideros cervinus*; time and frequency axes for each pulse have been adjusted to be equivalent).

**Appendix S3**

Table S3.1: Regional data on elevational distribution of bats after Bonaccorso (1998; with reference to updated nomenclature below the table).

| Family | Latin | 200 | 700 | 1200 | 1700 | 2200 | 2700 | 3200 | 3700 |
| --- | --- | --- | --- | --- | --- | --- | --- | --- | --- |
| **PTEROPODIDAE** | *Pteropus conspicillatus* | 1 | 0 | 0 | 0 | 0 | 0 | 0 | 0 |
| **EMBALLONURIDAE** | *Saccolaimus saccolaimus* | 1 | 0 | 0 | 0 | 0 | 0 | 0 | 0 |
| **VESPERTILIONIDAE** | *Pipistrellus wattsi* | 1 | 0 | 0 | 0 | 0 | 0 | 0 | 0 |
| **HIPPOSIDERIDAE** | *Hipposideros maggietaylorae* | 1 | 0 | 0 | 0 | 0 | 0 | 0 | 0 |
| **VESPERTILIONIDAE** | *Chalinolobus nigrogriseus* | 1 | 0 | 0 | 0 | 0 | 0 | 0 | 0 |
| **MOLOSSIDAE** | *Mormopterus beccarii^1^* | 1 | 0 | 0 | 0 | 0 | 0 | 0 | 0 |
| **MOLOSSIDAE** | *Otomops papuensis* | 1 | 0 | 0 | 0 | 0 | 0 | 0 | 0 |
| **PTEROPODIDAE** | *Pteropus hypomelanus* | 1 | 0 | 0 | 0 | 0 | 0 | 0 | 0 |
| **VESPERTILIONIDAE** | *Nyctophilus bifax^2^* | 1 | 0 | 0 | 0 | 0 | 0 | 0 | 0 |
| **PTEROPODIDAE** | *Pteropus macrotis* | 1 | 0 | 0 | 0 | 0 | 0 | 0 | 0 |
| **PTEROPODIDAE** | *Dobsonia minor* | 1 | 0 | 0 | 0 | 0 | 0 | 0 | 0 |
| **HIPPOSIDERIDAE** | *Aselliscus tricuspidatus* | 1 | 0 | 0 | 0 | 0 | 0 | 0 | 0 |
| **HIPPOSIDERIDAE** | *Hipposideros calcaratus* | 1 | 0 | 0 | 0 | 0 | 0 | 0 | 0 |
| **HIPPOSIDERIDAE** | *Hipposideros muscinus* | 1 | 0 | 0 | 0 | 0 | 0 | 0 | 0 |
| **PTEROPODIDAE** | *Nyctimene aello* | 1 | 1 | 0 | 0 | 0 | 0 | 0 | 0 |
| **PTEROPODIDAE** | *Paranyctimene raptor* | 1 | 1 | 1 | 0 | 0 | 0 | 0 | 0 |
| **PTEROPODIDAE** | *Macroglossus minimus* | 1 | 1 | 1 | 0 | 0 | 0 | 0 | 0 |
| **EMBALLONURIDAE** | *Emballonura furax* | 1 | 1 | 1 | 0 | 0 | 0 | 0 | 0 |
| **VESPERTILIONIDAE** | *Myotis moluccarum* | 1 | 1 | 1 | 0 | 0 | 0 | 0 | 0 |
| **HIPPOSIDERIDAE** | *Hipposideros diadema* | 1 | 1 | 1 | 0 | 0 | 0 | 0 | 0 |
| **VESPERTILIONIDAE** | *Pipistrellus papuanus* | 1 | 1 | 1 | 0 | 0 | 0 | 0 | 0 |
| **VESPERTILIONIDAE** | *Phoniscus papuensis* | 1 | 1 | 1 | 0 | 0 | 0 | 0 | 0 |
| **PTEROPODIDAE** | *Pteropus neohibernicus* | 1 | 1 | 1 | 0 | 0 | 0 | 0 | 0 |
| **EMBALLONURIDAE** | *Emballonura dianae* | 1 | 1 | 1 | 0 | 0 | 0 | 0 | 0 |
| **HIPPOSIDERIDAE** | *Hipposideros cervinus* | 1 | 1 | 1 | 0 | 0 | 0 | 0 | 0 |
| **MOLOSSIDAE** | *Chaerephon jobensis^3^* | 1 | 1 | 1 | 0 | 0 | 0 | 0 | 0 |
| **VESPERTILIONIDAE** | *Nyctophilus microtis* | 1 | 1 | 1 | 0 | 0 | 0 | 0 | 0 |
| **EMBALLONURIDAE** | *Emballonura beccarii* | 1 | 1 | 1 | 0 | 0 | 0 | 0 | 0 |
| **VESPERTILIONIDAE** | *Miniopterus australis* | 1 | 1 | 1 | 0 | 0 | 0 | 0 | 0 |
| **EMBALLONURIDAE** | *Emballonura raffrayana* | 1 | 1 | 1 | 0 | 0 | 0 | 0 | 0 |
| **EMBALLONURIDAE** | *Mosia nigrescens* | 1 | 1 | 1 | 0 | 0 | 0 | 0 | 0 |
| **RHINOLOPHIDAE** | *Rhinolophus megaphyllus* | 1 | 1 | 1 | 0 | 0 | 0 | 0 | 0 |
| **VESPERTILIONIDAE** | *Nyctophilus timoriensi^4^* | 1 | 1 | 1 | 0 | 0 | 0 | 0 | 0 |
| **VESPERTILIONIDAE** | *Kerivoula muscina* | 1 | 1 | 1 | 0 | 0 | 0 | 0 | 0 |
| **VESPERTILIONIDAE** | *Miniopterus propitristis^5^* | 1 | 1 | 1 | 0 | 0 | 0 | 0 | 0 |
| **PTEROPODIDAE** | *Nyctimene albiventer* | 1 | 1 | 1 | 1 | 0 | 0 | 0 | 0 |
| **HIPPOSIDERIDAE** | *Hipposideros ater* | 1 | 1 | 1 | 1 | 0 | 0 | 0 | 0 |
| **PTEROPODIDAE** | *Rousettus amplexicaudatus* | 1 | 1 | 1 | 1 | 0 | 0 | 0 | 0 |
| **RHINOLOPHIDAE** | *Rhinolophus euryotis* | 1 | 1 | 1 | 1 | 0 | 0 | 0 | 0 |
| **VESPERTILIONIDAE** | *Miniopterus magnater* | 1 | 1 | 1 | 1 | 0 | 0 | 0 | 0 |
| **MOLOSSIDAE** | *Otomops secundus* | 1 | 1 | 1 | 1 | 0 | 0 | 0 | 0 |
| **VESPERTILIONIDAE** | *Philetor brachypterus* | 1 | 1 | 1 | 1 | 0 | 0 | 0 | 0 |
| **VESPERTILIONIDAE** | *Scotorepens sanborni* | 1 | 1 | 1 | 1 | 1 | 0 | 0 | 0 |
| **VESPERTILIONIDAE** | *Pipistrellus angulatus* | 1 | 1 | 1 | 1 | 1 | 0 | 0 | 0 |
| **VESPERTILIONIDAE** | *Miniopterus medius* | 1 | 1 | 1 | 1 | 1 | 0 | 0 | 0 |
| **PTEROPODIDAE** | *Dobsonia moluccensis^6^* | 1 | 1 | 1 | 1 | 1 | 1 | 0 | 0 |
| **VESPERTILIONIDAE** | *Miniopterus schreibersii^7^* | 1 | 1 | 1 | 1 | 1 | 1 | 0 | 0 |
| **PTEROPODIDAE** | *Syconycteris australis* | 1 | 1 | 1 | 1 | 1 | 1 | 0 | 0 |
| **VESPERTILIONIDAE** | *Miniopterus macrocneme* | 1 | 1 | 1 | 1 | 1 | 1 | 1 | 0 |
| **HIPPOSIDERIDAE** | *Hipposideros edwardshilli* | 1 | 0 | 0 | 0 | 0 | 0 | 0 | 0 |
| **HIPPOSIDERIDAE** | *Hipposideros wollastoni* | 0 | 1 | 1 | 1 | 1 | 0 | 0 | 0 |
| **VESPERTILIONIDAE** | *Murina florium* | 0 | 1 | 1 | 1 | 1 | 1 | 0 | 0 |
| **RHINOLOPHIDAE** | *Rhinolophus arcuatus^8^* | 0 | 1 | 1 | 0 | 0 | 0 | 0 | 0 |
| **HIPPOSIDERIDAE** | *Hipposideros semoni* | 0 | 1 | 1 | 0 | 0 | 0 | 0 | 0 |
| **VESPERTILIONIDAE** | *Pipistrellus collinus* | 0 | 1 | 1 | 1 | 1 | 1 | 0 | 0 |
| **PTEROPODIDAE** | *Nyctimene cyclotis^9^* | 0 | 0 | 1 | 1 | 1 | 0 | 0 | 0 |
| **RHINOLOPHIDAE** | *Rhinolophus philippinensis* | 0 | 0 | 1 | 0 | 0 | 0 | 0 | 0 |
| **PTEROPODIDAE** | *Aproteles bulmerae* | 0 | 0 | 1 | 1 | 1 | 0 | 0 | 0 |
| **HIPPOSIDERIDAE** | *Hipposideros corynophyllus* | 0 | 0 | 1 | 1 | 0 | 0 | 0 | 0 |
| **VESPERTILIONIDAE** | *Nyctophilus microdon* | 0 | 0 | 1 | 0 | 1 | 0 | 0 | 0 |
| **MOLOSSIDAE** | *Tadarida kuboriensis^10^* | 0 | 0 | 1 | 0 | 1 | 1 | 0 | 0 |
| **PTEROPODIDAE** | *Syconycteris hobbit* | 0 | 0 | 1 | 0 | 1 | 0 | 0 | 0 |

*^1^Ozimops beccarii* (Jackson and Groves 2015)

*^2^Nyctophilus bifax is no longer in PNG* (Parnaby 2009)

*^3^Mops jobensis* (Gregorin & Cirranello 2016)

*^4^Nyctophilus shirleyae* (Parnaby 2009)

*^5^Miniopterus protristis* (IUCN 2021)

*^6^Dobsonia magna* (IUCN 2021)

*^7^Miniopterus orianae* (Jackson and Groves 2015)

*^8^Rhinolophus mcintyrei* (Patrick et al. 2013)

*^9^Nyctimene certans* (Irwin 2017)

*^10^Austronomus kuboriensis* (Gregorin & Cirranello 2016)

Table S3.2: Explanatory variables and GPS location at each site along the Mt. Wilhelm. F.tree ab= Fruiting tree abundance; F.tree sp= Fruiting tree richness; Moth ab= Moth abundance; Moth sp= Moth richness.

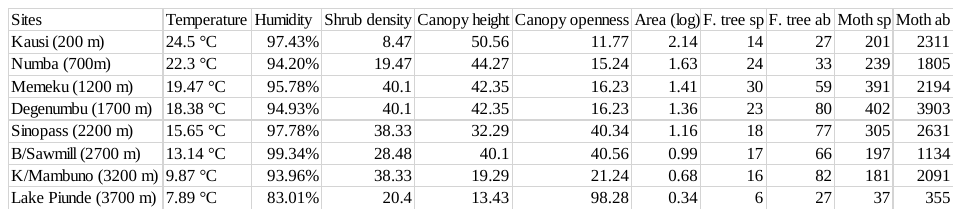

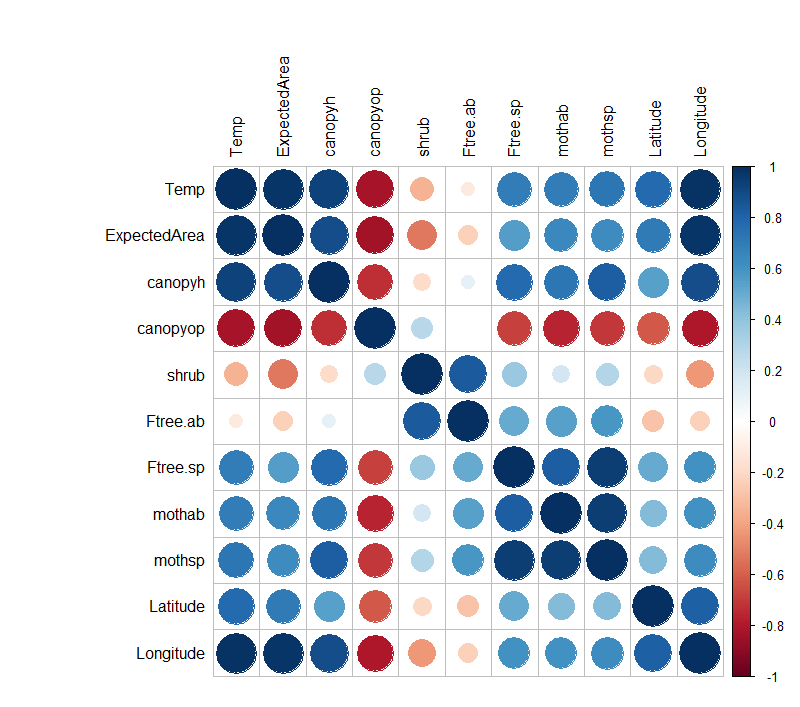

Fig. S3.1: Correlation table of all the explanatory variables used in the models. Dark blue means a very high positive correlation between two variables. Temp=Temperature; Canopyh= Canopy height; Canopyop= Canopy openness; Shrub= Shrub density; Ftree.ab= Fruiting tree abundance; Ftree.sp= Fruiting tree richness; Mothab= Moth abundance; Mothsp= Moth richness.

**Appendix S4**

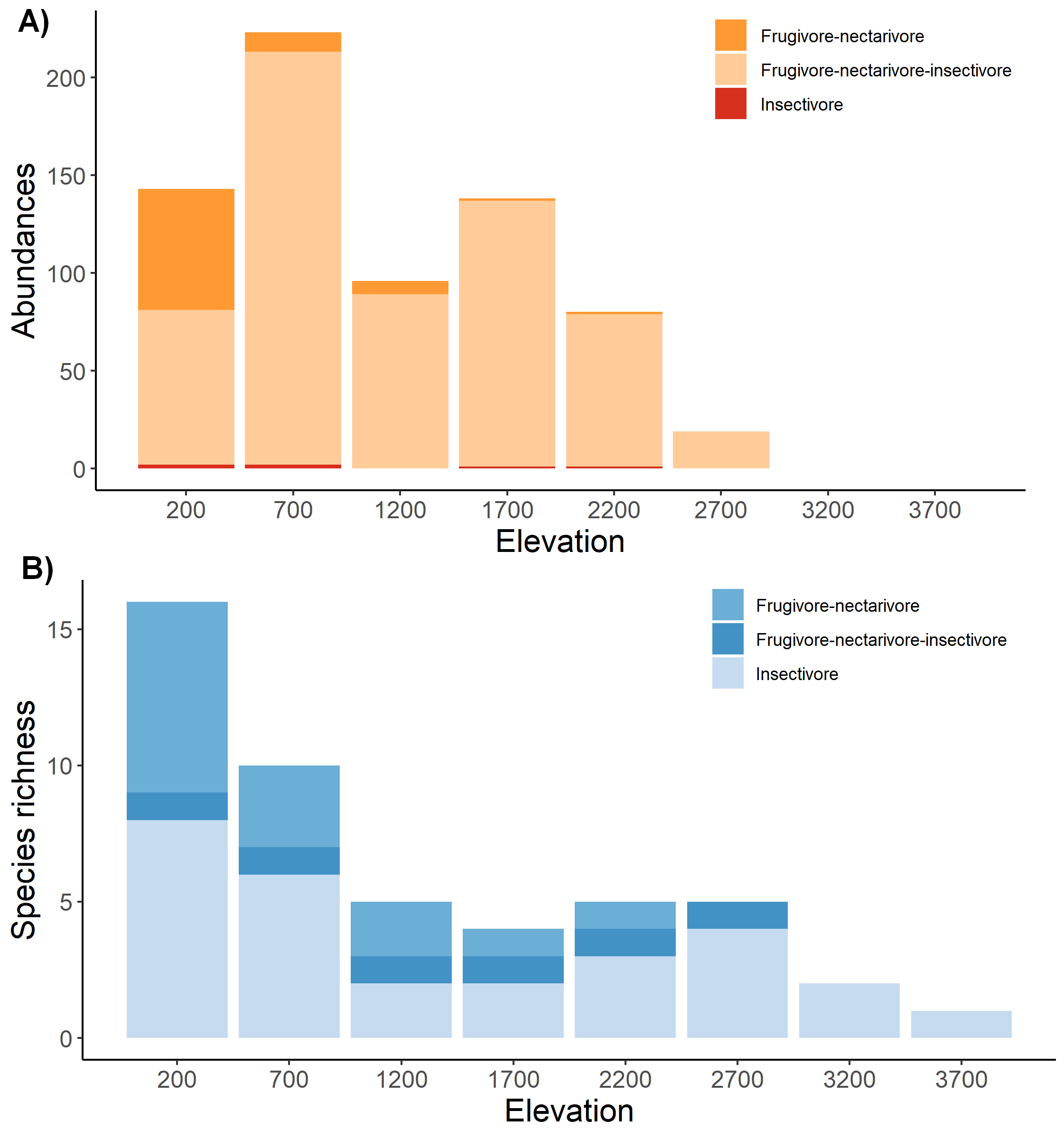

Fig. S4.1: A) Bar chart representing bat abundances partitioned in three feeding guilds along the elevational gradient of Mt. Wilhelm from mist-netting data. B) Bar chart of the total species richness partitioned in three feeding guilds along the elevational gradient of Mt. Wilhelm from acoustic and mist-netting data.

**Appendix S5**

Table S5.1: Akaike's second-order information criterion (AIC_c_) for models of observed species richness along the elevational gradient, estimated for all bat observations and for the same observations divided amongst two feeding guilds. Models comprise all combinations of five or four explanatory variables.

| *All bats* | | | |  |  |  |  |  |  |  |  |  |  |  |  |  |  |
| --- | --- | --- | --- | --- | --- | --- | --- | --- | --- | --- | --- | --- | --- | --- | --- | --- | --- |
| Temperature | Area | | Canopy open. | | Canopy height | Shrub dens. | | df | | Log (L) | | | AICc | | ΔAICc | | W_1_ |
|  | **1.31** | |  | |  |  | | 2.00 | | -14.55 | | | 35.50 | | 0.00 | | 0.46 |
| 2.12 |  | |  | |  |  | | 2.00 | | -15.32 | | | 37.00 | | 1.54 | | 0.21 |
|  |  | |  | | 4.72 |  | | 2.00 | | -16.31 | | | 39.00 | | 3.52 | | 0.08 |
|  |  | |  | | 3.31 | -1.22 | | 3.00 | | -14.26 | | | 40.50 | | 5.01 | | 0.04 |
| 1.58 |  | |  | |  | -0.92 | | 3.00 | | -14.31 | | | 40.60 | | 5.12 | | 0.04 |
|  | 1.59 | | 0.70 | |  |  | | 3.00 | | -14.38 | | | 40.80 | | 5.26 | | 0.03 |
|  | 1.07 | |  | | 1.10 |  | | 3.00 | | -14.43 | | | 40.90 | | 5.35 | | 0.03 |
|  | 1.22 | |  | |  | -0.20 | | 3.00 | | -14.53 | | | 41.10 | | 5.55 | | 0.03 |
| 0.29 | 1.15 | |  | |  |  | | 3.00 | | -14.54 | | | 41.10 | | 5.56 | | 0.03 |
| 2.57 |  | | 0.65 | |  |  | | 3.00 | | -15.21 | | | 42.40 | | 6.91 | | 0.01 |
| 1.70 |  | |  | | 1.07 |  | | 3.00 | | -15.26 | | | 42.50 | | 7.01 | | 0.01 |
|  |  | |  | |  | -2.02 | | 2.00 | | -18.80 | | | 44.00 | | 8.49 | | 0.01 |
|  |  | | -2.19 | |  |  | | 2.00 | | -18.86 | | | 44.10 | | 8.61 | | 0.01 |
|  |  | | -0.65 | | 3.84 |  | | 3.00 | | -16.07 | | | 44.10 | | 8.64 | | 0.01 |
|  |  | | -1.42 | |  | -1.38 | | 3.00 | | -16.11 | | | 44.20 | | 8.72 | | 0.01 |
| 0.76 |  | |  | | 1.92 | -1.03 | | 4.00 | | -14.08 | | | 49.50 | | 13.99 | | 0.00 |
|  | 0.42 | |  | | 2.33 | -0.82 | | 4.00 | | -14.17 | | | 49.70 | | 14.16 | | 0.00 |
|  | 1.36 | | 0.85 | | 1.35 |  | | 4.00 | | -14.20 | | | 49.70 | | 14.23 | | 0.00 |
| 2.04 |  | | 0.64 | |  | -0.92 | | 4.00 | | -14.20 | | | 49.70 | | 14.23 | | 0.00 |
|  |  | | -0.11 | | 3.17 | -1.19 | | 4.00 | | -14.25 | | | 49.80 | | 14.33 | | 0.00 |
| 0.75 | 1.27 | | 0.94 | |  |  | | 4.00 | | -14.29 | | | 49.90 | | 14.40 | | 0.00 |
| 1.73 | -0.12 | |  | |  | -1.00 | | 4.00 | | -14.31 | | | 50.00 | | 14.45 | | 0.00 |
|  | 1.64 | | 0.75 | |  | 0.07 | | 4.00 | | -14.38 | | | 50.10 | | 14.59 | | 0.00 |
| -0.36 | 1.20 | |  | | 1.42 |  | | 4.00 | | -14.41 | | | 50.20 | | 14.65 | | 0.00 |
| 2.19 |  | | 0.63 | | 0.97 |  | | 4.00 | | -15.15 | | | 51.60 | | 16.14 | | 0.00 |
|  |  | |  | |  |  | | 1.00 | | -25.01 | | | 52.70 | | 17.19 | | 0.00 |
| 1.22 |  | | 0.57 | | 1.77 | -1.02 | | 5.00 | | -14.00 | | | 68.00 | | 32.49 | | 0.00 |
| 1.30 | -0.49 | |  | | 2.05 | -1.35 | | 5.00 | | -14.06 | | | 68.10 | | 32.60 | | 0.00 |
|  | 0.80 | | 0.51 | | 2.08 | -0.56 | | 5.00 | | -14.11 | | | 68.20 | | 32.71 | | 0.00 |
| 1.68 | 0.33 | | 0.72 | |  | -0.71 | | 5.00 | | -14.19 | | | 68.40 | | 32.87 | | 0.00 |
| 0.19 | 1.31 | | 0.90 | | 1.21 |  | | 5.00 | | -14.20 | | | 68.40 | | 32.89 | | 0.00 |
| 1.33 | -0.12 | | 0.53 | | 1.82 | -1.10 | | 6.00 | | -14.00 | | | 124.00 | | 88.49 | | 0.00 |
| *Frugivore-nectarivore* | | | |  |  |  |  |  |  |  |  |  |  |  |  |  |  |
| Temperature | | Area | Tree fr. abun. | | Tree fr. sp. | df | Log (L) | | | | | AICc | | ΔAICc | | W_1_ | |
| 3.62 | |  |  | |  | 2.00 | -9.31 | | | | | 25.00 | | 0.00 | | 0.55 | |
|  | | 1.79 |  | |  | 2.00 | -9.96 | | | | | 26.30 | | 1.29 | | 0.29 | |
| 3.95 | |  | 0.19 | |  | 3.00 | -9.29 | | | | | 30.60 | | 5.55 | | 0.03 | |
| 3.19 | | 0.23 |  | |  | 3.00 | -9.30 | | | | | 30.60 | | 5.57 | | 0.03 | |
| 3.59 | |  |  | | -0.12 | 3.00 | -9.30 | | | | | 30.60 | | 5.58 | | 0.03 | |
|  | | 2.07 |  | | 0.91 | 3.00 | -9.39 | | | | | 30.80 | | 5.76 | | 0.03 | |
|  | | 2.25 | 0.55 | |  | 3.00 | -9.79 | | | | | 31.60 | | 6.55 | | 0.02 | |
|  | |  | -1.64 | | 1.44 | 3.00 | -12.51 | | | | | 37.00 | | 12.00 | | 0.00 | |
|  | |  |  | |  | 1.00 | -17.45 | | | | | 37.60 | | 12.53 | | 0.00 | |
|  | |  | -0.93 | |  | 2.00 | -15.65 | | | | | 37.70 | | 12.67 | | 0.00 | |
|  | |  |  | | 0.77 | 2.00 | -16.55 | | | | | 39.50 | | 14.46 | | 0.00 | |
| 4.29 | |  | 0.45 | | -0.39 | 4.00 | -9.22 | | | | | 39.80 | | 14.74 | | 0.00 | |
| 3.15 | | 0.62 | 0.39 | |  | 4.00 | -9.23 | | | | | 39.80 | | 14.76 | | 0.00 | |
| 3.18 | | 0.24 |  | | 0.00 | 4.00 | -9.30 | | | | | 39.90 | | 14.91 | | 0.00 | |
|  | | 2.31 | 0.28 | | 0.86 | 4.00 | -9.36 | | | | | 40.10 | | 15.03 | | 0.00 | |
| 4.37 | | 0.05 | 0.45 | | -0.41 | 5.00 | -9.22 | | | | | 58.40 | | 33.41 | | 0.00 | |
| *Insectivore* | | | |  |  |  |  |  |  |  |  |  |  |  |  |  |  |
| Temperature | | Area | Moth abun. | | Moth sp. | df | | | Log (L) | | AICc | | | | ΔAICc | | W_1_ |
|  | | **0.98** |  | |  | 2.00 | | | -13.99 | | 34.40 | | | | 0.00 | | 0.48 |
| 1.48 | |  |  | |  | 2.00 | | | -14.43 | | 35.30 | | | | 0.87 | | 0.31 |
|  | |  |  | |  | 1.00 | | | -18.37 | | 39.40 | | | | 5.03 | | 0.04 |
|  | | 1.08 | -0.43 | |  | 3.00 | | | -13.88 | | 39.80 | | | | 5.37 | | 0.03 |
|  | | 1.00 |  | | -0.15 | 3.00 | | | -13.97 | | 39.90 | | | | 5.57 | | 0.03 |
| -0.08 | | 1.03 |  | |  | 3.00 | | | -13.99 | | 40.00 | | | | 5.60 | | 0.03 |
| 1.74 | |  |  | | -0.70 | 3.00 | | | -14.12 | | 40.20 | | | | 5.85 | | 0.03 |
| 1.77 | |  | -0.71 | |  | 3.00 | | | -14.16 | | 40.30 | | | | 5.95 | | 0.03 |
|  | |  | 0.76 | |  | 2.00 | | | -17.67 | | 41.70 | | | | 7.36 | | 0.01 |
|  | |  |  | | 0.70 | 2.00 | | | -17.74 | | 41.90 | | | | 7.50 | | 0.01 |
|  | |  | 0.61 | | 0.17 | 3.00 | | | -17.67 | | 47.30 | | | | 12.95 | | 0.00 |
|  | | 1.13 | -1.10 | | 0.68 | 4.00 | | | -13.78 | | 48.90 | | | | 14.51 | | 0.00 |
| 0.35 | | 0.89 | -0.51 | |  | 4.00 | | | -13.86 | | 49.10 | | | | 14.67 | | 0.00 |
| 0.35 | | 0.81 |  | | -0.27 | 4.00 | | | -13.97 | | 49.30 | | | | 14.88 | | 0.00 |
| 1.77 | |  | -0.25 | | -0.51 | 4.00 | | | -14.11 | | 49.50 | | | | 15.16 | | 0.00 |
| -0.83 | | 1.63 | -1.41 | | 1.20 | 5.00 | | | -13.75 | | 67.50 | | | | 33.11 | | 0.00 |

**Appendix S6**

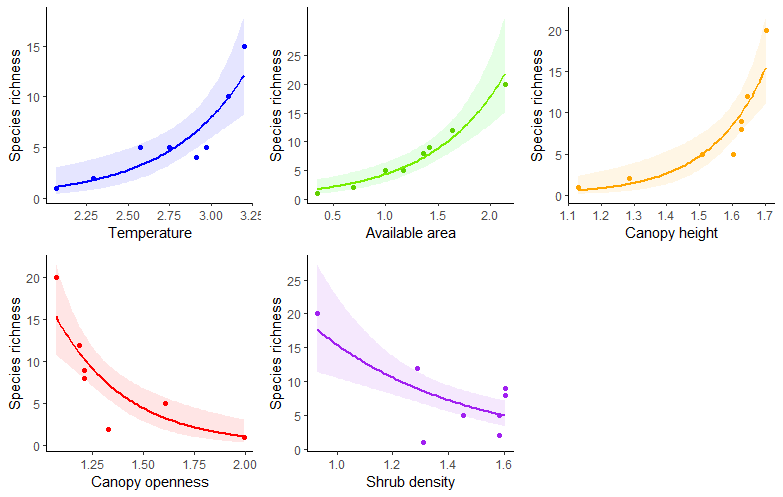

Fig. S6.1: Fitted models of the total species richness according to five explanatory variables: mean daily temperature, available land area, canopy height, canopy openness and shrub density.

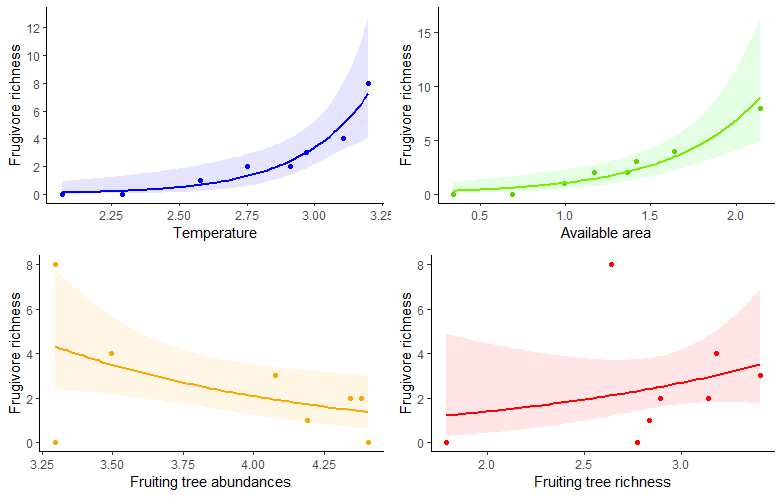

Fig. S6.2: Fitted models of the frugivore-nectarivore species richness according to four explanatory variables: mean daily temperature, available land area, fruiting tree richness and abundance.

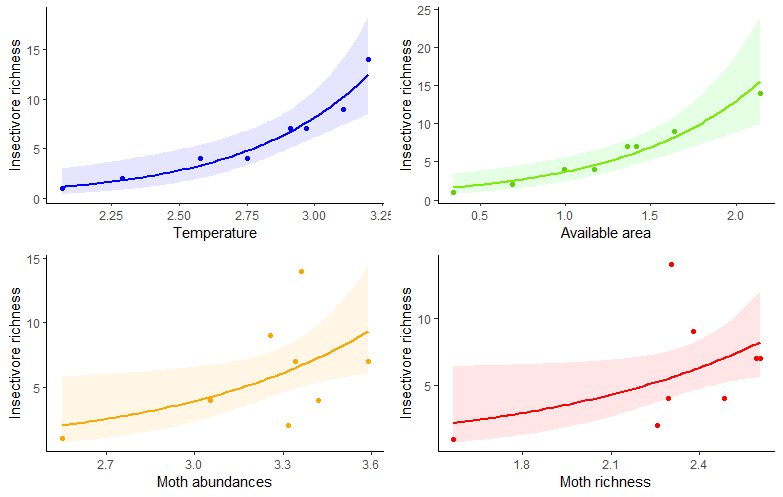

Fig. S6.3: Fitted models of the insectivore species richness according to four explanatory variables: mean daily temperature, available land area, moth abundance and richness.

**Appendix S7**

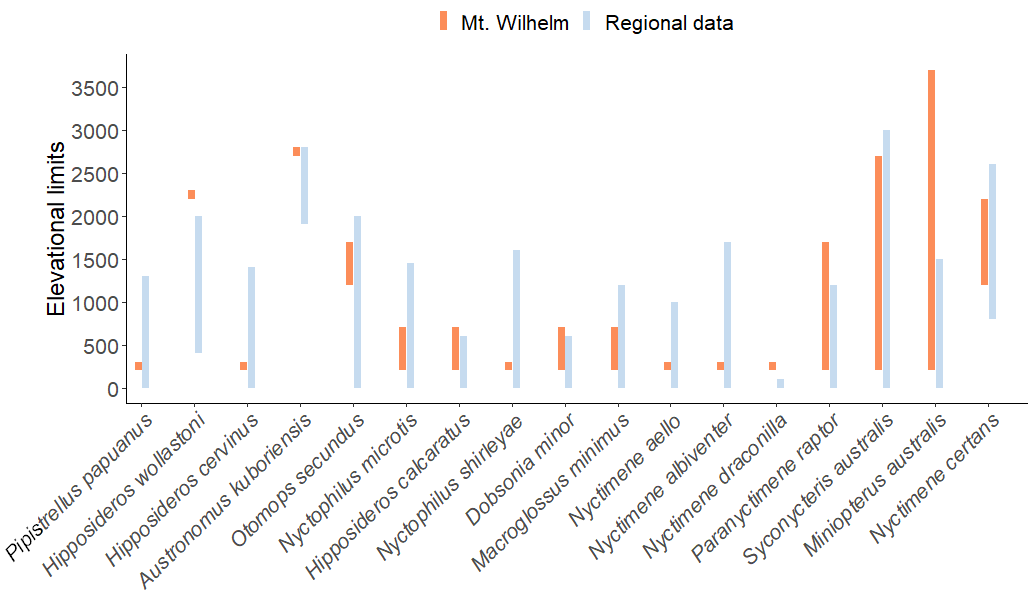

Fig. S7.1: Comparison of the elevational ranges of the bat species found in Mt. Wilhelm with regional data (Bonaccorso 1998). Note that calls attributed to none or more than one species are not presented here.

**Appendix S8**

Table S8.1: Roosting and foraging information found in Bonaccorso (1998; using current nomenclature) for the species detected in Mt. Wilhelm.

| **Species** | **Caves** | **Trees** | **Buildings** | **Foraging** | **Diet** |
| --- | --- | --- | --- | --- | --- |
| **HIPPOSIDERIDAE** | | | | | |
| *Hipposideros wollastoni* | x |  |  | NA | NA |
| *Hipposideros cervinus* | x | x | x | Gleaner and aerial in dense forest and urban areas | Beetles, moths |
| *Hipposideros calcaratus* | x |  |  | NA | NA |
| **EMBALLONURIDAE** | | | | | |
| *Emballonura beccarii* | x |  |  | Gleaner, clear or dense forest, next to streams | Beetles |
| *Mosia nigrescens* | x | x | x | From canopy to ground level from forest to urban areas | Aerial and foliage-clinging insects, ants |
| **VESPERTILIONIDAE** | | | | | |
| *Pipistrellus papuanus* |  | x | x | Urban and open areas | Aerial insects |
| *Pipistrellus collinus* | NA | NA | NA | Native gardens | Aerial insects |
| *Nyctophilus microtis* | x | x |  | NA | NA |
| *Nyctophilus shirleyae* |  | x |  | Gleaner, subcanopy and over water | Moths and beetles |
| **MINIOPTERIDAE** | | | | | |
| *Miniopterus tristis* | *x* |  |  | Above canopy, clear areas | Aerial insects |
| *Miniopterus australis* | x | x |  | Beneath the canopy | Aerial insects |
| **MOLOSSIDAE** | | | | | |
| *Austronomus kuboriensis* | NA | NA | NA | Hawking above canopy | Beetles |
| *Otomops secundus* | NA | NA | NA | Open areas, canopy and urban areas | Large insects, beetles |
| **PTEROPODIDAE** | | | | | |
| *Syconycteris australis* |  | x |  | Generalist | Moraceae, Piperaceae, Solanaceae, flowers, insects |
| *Macroglossus minimus* |  | x | x | Generalist | Pollen, nectar |
| *Paranyctimene raptor* |  | x |  | Gardens, swamps | Ficus, Piper |
| *Nyctimene albiventer* |  | x |  | Ground or around trees | NA |
| *Nyctimene draconilla* |  | x |  | Fresh swamp, river | Fruits |
| *Nyctimene aello* |  | x |  | Subcanopy | Figs |
| *Nyctimene certans* |  | x |  | Ground and subcanopy | Maybe figs |
| *Dobsonia minor* |  | x |  | Dense understory | Figs, piper |
